# Establishment of titration-based control of DNA replication in *Escherichia coli*

**DOI:** 10.64898/2026.05.06.723188

**Authors:** Belén Adiego-Pérez, Demian Fluit, Christina Ludwig, Mareike Berger, Johannes Hohlbein, Raymond H.J. Staals, Pieter Rein ten Wolde, John van der Oost, Nico J. Claassens, Lorenzo Olivi

**Affiliations:** Laboratory of Microbiology, Wageningen University & Research, Wageningen, the Netherlands; Bavarian Center for Biomolecular Mass Spectrometry (BayBioMS), School of Life Sciences, Technical University of Munich, Germany; Institute AMOLF, Amsterdam, the Netherlands; Laboratory of Biophysics, Wageningen University & Research, Wageningen, the Netherlands; Microspectroscopy Research Facility, Wageningen University & Research, Wageningen, the Netherlands

## Abstract

*Escherichia coli* couples the initiation of DNA replication with cell size by modulating the activity of the replication initiator protein DnaA. The activity of DnaA is regulated by both its interconversion between an active and inactive form and its titration on binding sites on the chromosome. Whereas its interconversion has been thoroughly studied, the extent to which DnaA titration can control replication initiation is poorly understood. Here, we describe the control of *E. coli* DNA replication via titration by modulating the expression of an ‘always*-*active’ DnaA variant in four growth conditions. While we obtained stable cell cycles during slow growth, faster growth associated with overlapping replication forks led to replicative instability and DNA damage. Overall, our results provide insights into the limits of titration-based systems in the control of genome replication and their potential role in the evolutionary trajectory of *E. coli*. Finally, this study provides design principles for a simplified, titration-only regulatory mechanism for DNA replication in synthetic cells.

## Introduction

Precise control over genome replication is crucial for optimal fitness of all forms of life. In most studied bacteria, the initiation of DNA replication relies on the replication initiator protein DnaA^1^. The binding of DnaA to specific DNA motifs (DnaA boxes) present at the origin of replication of the bacterial chromosome causes local melting of the double-stranded DNA structure and recruitment of the proteins of the replisome complex^2,3^. Uncontrolled initiation of DNA replication is associated with hyper-replicative stress and generally leads to severe fitness loss^4,5^. Thus, several mechanisms are in place to ensure that DnaA binds and unwinds the origin only when specific conditions are met.

In *Escherichia coli*, cells coordinate DNA replication with growth^6,7^, such that initiation occurs at a constant cellular volume per number of origins^7,8^. At fast growth, this coupling results in the presence of multiple replication forks in a single chromosome, which allows cells to divide faster than the time needed to replicate their chromosome^9,10^. The known mechanisms controlling this characteristic behaviour are the regulation of both production and activity of the DnaA protein. First, the near-balanced expression of the *dnaA* gene results in constant intracellular DnaA concentrations^11^, with the specific concentration value set by the growth rate^12^. Second, DnaA exists either in an active or an inactive form, bound to ATP or ADP, respectively^13^. The ratio of the two forms is controlled by several mechanisms, involving the interaction of DnaA with either non-coding chromosomal loci^14,15^, the regulatory protein Hda^16^, or the cell membrane^17^. Only the active, ATP-bound form can unwind the origin of replication (*oriC*)^18–20^. Thus, accumulation of DnaA-ATP to a certain threshold value is a critical step in the initiation of DNA replication^21^. At the same time, the titration of DnaA on DNA boxes spread across the chromosome of *E. coli* also acts in regulating replication initiation^22,23^. In this titration model, the free concentration of DnaA is maintained at a low level through sequestration of the protein on high-affinity DnaA boxes spread across the chromosome. Once these titration sites are saturated, the free concentration of DnaA rises until low affinity DnaA boxes present on *oriC* become significantly occupied^18^, eventually resulting in initiation of replication. Although evidence on the effect of titration has historically been scarce, recent studies support a scenario where a combination of both mechanisms orchestrates cell cycles in *E. coli*. Previously, we probed the mobility of DnaA in live *E. coli* cells and demonstrated that its titration indeed plays a role in the control of DNA replication during slow growth^24^. Cell cycle simulations also predicted that only the combination of titration and interconversion of DnaA yields stable replication cycles across growth conditions^25,26^. Finally, additional indirect evidence comes from the fact that reaching peak levels of DnaA-ATP is necessary but not sufficient to initiate DNA replication^27^, hinting at DnaA titration being responsible for the observed delay.

To date, the extent to which DnaA titration alone can contribute to the regulation of DNA replication is still unclear. Here, we used a homeostatic, inducible genetic circuit to control the expression of the mutant DnaA cold-sensitive allele (DnaAcos), a variant impaired in nucleotide binding^28^. This modification abolishes interconversion between active and inactive DnaA and leaves titration as the only remaining, known control mechanism for DNA replication. This genetic system allowed us to experimentally confirm predictions of DNA replication instability in *E. coli* in the absence of an ATP/ADP switch in regimes of overlapping replication forks^25^, which should lead to an unviable phenotype^4^. Surprisingly, the selection of a missense mutation in the DnaB helicase gene restored growth in all the tested conditions. We thus analysed the phenotype of the resulting *E. coli* Co-Regulatory TetR-DnaAcos (*E. coli* CoRTeDcos) strain in different growth conditions and induction levels by measuring a variety of cellular parameters in parallel. We conclude that a titration-based system recreates the phenomenon of constant volumes of initiation across growth conditions, and that this volume is uniquely set by the concentration of DnaAcos. Nevertheless, proteomic analysis revealed signs of hyper‐replicative stress in *E. coli* CoRTeDcos under conditions that promote overlapping replication forks. Altogether, our results bring new perspectives on DnaA titration as a form of control of DNA replication initiation and provide valuable insights for the eventual design of controlled genetic inheritance systems in synthetic cells.

## Results

### Construction of an *E. coli* strain incapable of DnaA-ATP/DnaA-ADP switch

In this study, we sought to create a molecular mechanism based on titration that recreates the phenomenological observation of a constant initiation volume across growth rates. To achieve this behaviour, several design criteria needed to be met. First, the native switch between DnaA-ATP and DnaA-ADP had to be abolished. An *E. coli* strain with deletions for of all known loci involved in the interconversion between the two forms has already been reported^11^. Yet, this strain was still capable of cycling between the two forms via the intrinsic ATPase activity of DnaA and the binding of ATP upon *de novo* synthesis^11^. Thus, we decided to replace the wild-type DnaA with the mutant allele DnaAcos. DnaAcos contains two amino acid substitutions that impair nucleotide binding (A184V and H252Y)^28,29^ and two additional substitutions (Q156L and Y271H) that stabilise its activity below 42 °C^28,30^. As a consequence, DnaAcos is constitutively active in initiating DNA replication^28^. Next, in pure titration-based systems, the rate at which DnaA is synthesised dictates both the mass fraction of DnaA (*ϕ*_DnaA_) and how quickly the DnaA boxes on the chromosome are saturated. Since the saturation of DnaA boxes leads to *oriC* binding and initiation of DNA replication, higher *ϕ*_DnaA_ then result in smaller volume of initiations^25^. Thus, achieving constant volumes of initiation across growth rates means implementing an expression system that maintains *ϕ*_DnaA_ constant independently of nutrient conditions. In wild-type *E. coli* wild-type, *ϕ*_DnaA_ varies of ∼50% across growth rates^12^. To achieve constant and controllable *ϕ*_DnaA_ across growth rates, we selected a genetic circuit where the expression of DnaAcos is controlled by a negatively autoregulated TetR repressor^31^. This type of genetic architecture should provide stable mass fractions of the co-regulated protein (i.e., DnaAcos) across growth rates, with the added benefit of controlling such concentration via addition of the inducer molecule anhydrotetracycline (aTc)^31^. Finally, replication needs to be initiated when a critical threshold of DnaAcos proteins has accumulated. In DnaA titration-based system, this threshold is set by the number of high-affinity DnaA boxes that need to be saturated before DnaA can bind and unwind *oriC*^22,25^. Previously, we showed that the chromosome of *E. coli* is capable of titrating DnaA^24^. As such, we chose to rely on the hundreds of native DnaA boxes distributed across the *E. coli* chromosome to govern the accumulation of DnaA toward its critical threshold.

Following these design principles, we replaced the wild-type *E. coli dnaA* gene and its promoters with a co-regulatory construct composed of the *P*_*LtetO-1*_ promoter, the *tetR* repressor gene and the *dnaAcos* gene (Figure 1A). SeqA-mediated repression of DnaAcos was also abolished by removing GATC motifs on the *P*_*dnaA*_ promoter^32,33^. Blockage of *oriC* during the eclipse period was maintained^32,33^. Notably, a co-regulatory construct yields low levels of protein production even in absence of the inducer^31^, ensuring DNA replication initiation and viability of our strain in absence of aTc. After successfully obtaining the mutant strain, we first characterised its growth phenotype in two growth media and five concentrations of aTc (Figure 1B, Figure S1). The obtained strain was viable in poor nutrient medium leading to slow growth in a variety of aTc concentrations, with growth rates slightly lower than the wild-type (*λ*_wt_ = 0.27 ± 0.015 h^-1^, *λ*_mutant_ = 0.207 ± 0.015 h^-1^). However, when cultivated in a medium leading to overlapping replication regime in the wild-type (*λ*_wt_ = 0.987 ± 0.016 h^-1^), the mutant strain was incapable to grow at concentrations of aTc of 10 ng/mL or higher. Notably, prolonged incubation in the same conditions yielded an evolved strain capable of proliferating at high growth rates and across a large range of aTc concentrations (Figure 1B).

**Figure 1.**
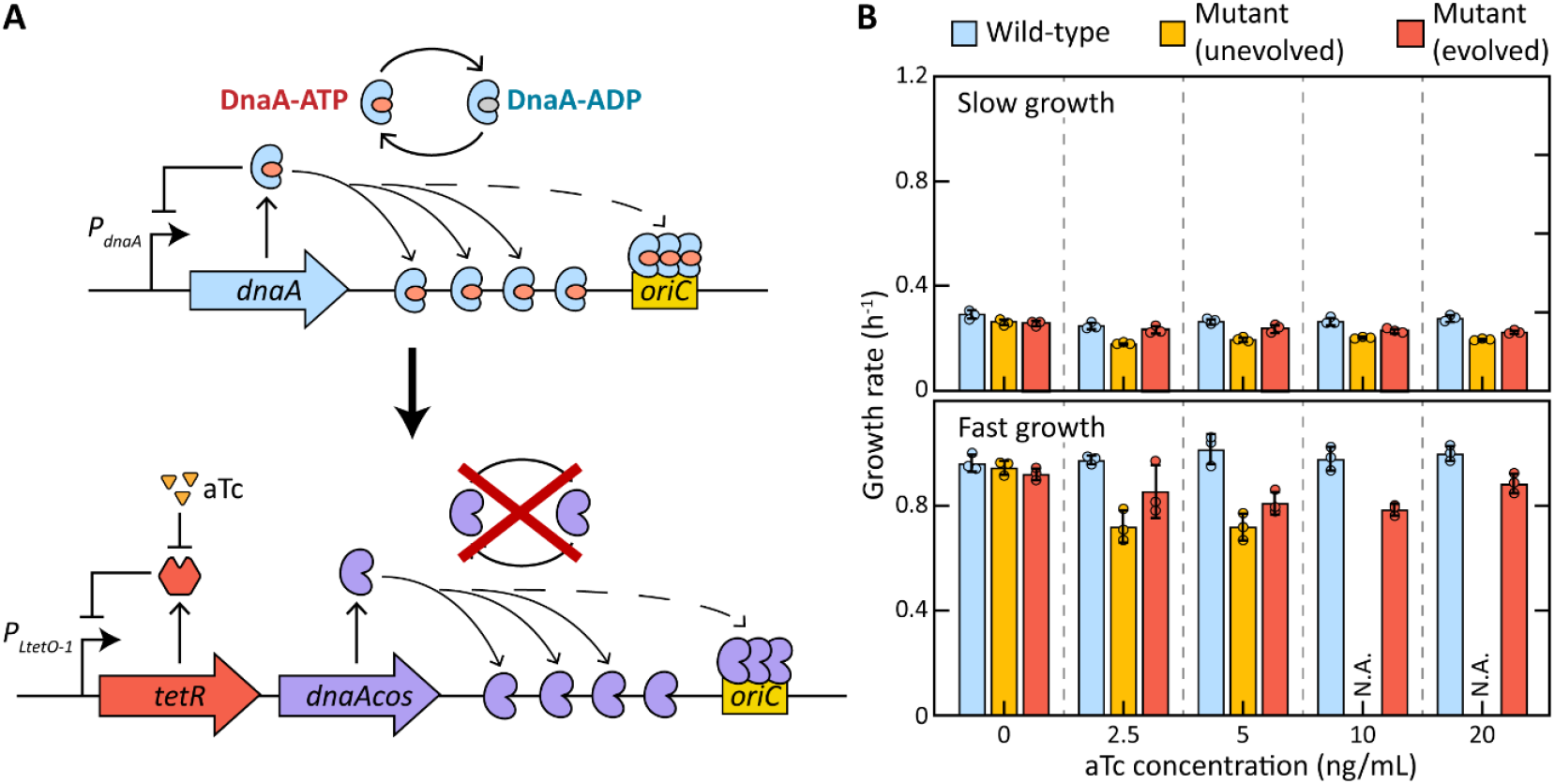
Design and characterisation of *E. coli* CoRTeDcos. **A)** Design of a genetic architecture allowing for titration-based control of DNA replication in *E. coli*. Top: native *dnaA* locus in *E. coli*, bottom: engineered locus in *E. coli* CoRTeDcos strain. **B)** Growth rate of the wild-type *E. coli* MG1655 (blue), the initial mutant strain (yellow) and the final *E. coli* CoRTeDcos strain (red) after the selection of the DnaB(M242I) substitution. Media leading to slow growth (top) or fast growth and overlapping replication forks (bottom) were tested in presence of varying concentrations of aTc. The final *E. coli* CoRTeDcos strain was selected during fast growth conditions and in presence of aTc. Growth curves are available in Supplementary Information (Figure S1).

Initiator titration systems are predicted to be unstable at growth rates leading to overlapping replication regimes^25,26^. These predictions are in line with the known hyper-replicative stress phenotype associated with DnaAcos^4,28,30^ and likely explains the lack of viability of the original mutant strain in fast growth conditions. Interestingly, the evolved strain is capable of surviving in the same conditions without any mutations in the co-regulatory *tetR*-*dnaAcos* locus, nor any previously reported mutation rescuing hyper-replicative phenotypes^34–36^. Rather, we observed a single mutation in the replicative DNA helicase *dnaB* gene, leading to an M242I amino acidic substitution. This mutation is known to cause several defects, resulting in a DNA helicase hexamer with reduced unwinding capacity^37^. Introduction of this mutation restored viability of the strain in a large range of growth conditions. We thus decided to further characterise this strain, which we named *E. coli* CoRTeDcos (Co-Regulatory TetR-DnaAcos). Notably, when first attempting to obtain the average number of origins of *E. coli* CoRTeDcos populations, we noticed that the strain seemed to be resistant to cell cycle synchronisation via rifampicin (Figure S2). Resistance to rifampicin treatment has already been described in *E. coli* strains where the levels of active DnaA remain high in the cell even after the stop of *dnaA* transcription, either by substantial overexpression of DnaA^38^ or via gene deletions (e.g. Δ*hda*^39,40^ and Δ*ihfAB*^39^). We reasoned that this similarity between *E. coli* CoRTeDcos and other hyper-replicating *E. coli* mutants serves as additional evidence of the constitutively active DnaAcos being the only replication initiator in the cell.

### Quantitative cell cycle relations in absence of DnaA-ATP/ADP interconversion

To assess the replicative behaviour of *E. coli* CoRTeDcos, we leveraged turbidostat cultivations to balance cells in exponential growth^7,12,24,41^ and performed population-level measurements of relevant cellular parameters^7^. We first benchmarked our strategy by reproducing known behaviours of wild-type *E. coli* across different nutrient conditions (Figure S3). We then moved to compare the behaviour of the wild type *E. coli* MG1655 and the mutant *E. coli* CoRTeDcos in four different media supporting growth at different growth rates (M1 to M4; see Table S2 for composition) and two levels of aTc (0 and 10 ng/mL) (Figure 2). In medium M1 and M2 (*λ*_WT-M1_ = 0.36 ± 0.03 h^-1^, *λ*_WT-M2_ = 0.64 ± 0.02 h^-1^), *E. coli* CoRTeDcos grew similarly to the wild-type at either 0 or 10 ng/mL of aTc (*λ*_CoRTeDcos-M1-0_ = 0.34 ± 0.03 h^-1^, *λ*_CoRTeDcos-M1-10_ = 0.38 ± 0.02 h^-1^; *λ*_CoRTeDcos-M2-0_ = 0.65 ± 0.02 h^-1^, *λ*_CoRTeDcos-M2-10_ = 0.67 ± 0.02 h^-1^). In more nutrient-rich conditions leading to overlapping replication forks, the presence of the mutant locus led to a reduction of growth rate. In medium M3, *E. coli* CoRTeDcos grew consistently slower than the wild-type, with the presence of aTc eliciting no additional effect (*λ*_WT_ = 1.25 ± 0.02 h^-1^, *λ*_CoRTeDcos-M3-0_ = 1.04 ± 0.01 h^-1^, *λ*_CoRTeDcos-M3-10_ = 1.08 ± 0.03 h^-1^). At even faster growth rates, (*λ*_WT_ = 1.83 ± 0.05 h^-1^), the presence of inducer seemed to further impact growth (*λ*_CoRTeDcos-0_ = 1.45 ± 0.03 h^-1^, *λ*_CoRTeDcos-10_ = 1.28 ± 0.08 h^-1^). To characterise the expression behaviour of the co-regulatory construct, we obtained *ϕ*_DnaA_ via quantitative proteomics (Figure 2B). In absence of inducer, the *ϕ*_DnaAcos_ doubled across growth rates, while remaining around 5.5 lower than the *ϕ*_DnaA_ in wild-type cells. In presence of 10 ng/mL of aTc, the co-regulatory construct yielded constant *ϕ*_DnaAcos_, varying by only 0.7% across 5-fold increases of growth rate. In contrast, the wild-type *ϕ*_DnaA_ varied of 43% in the same range of growth rates, similarly to what had previously been showed^12^.

**Figure 2.**
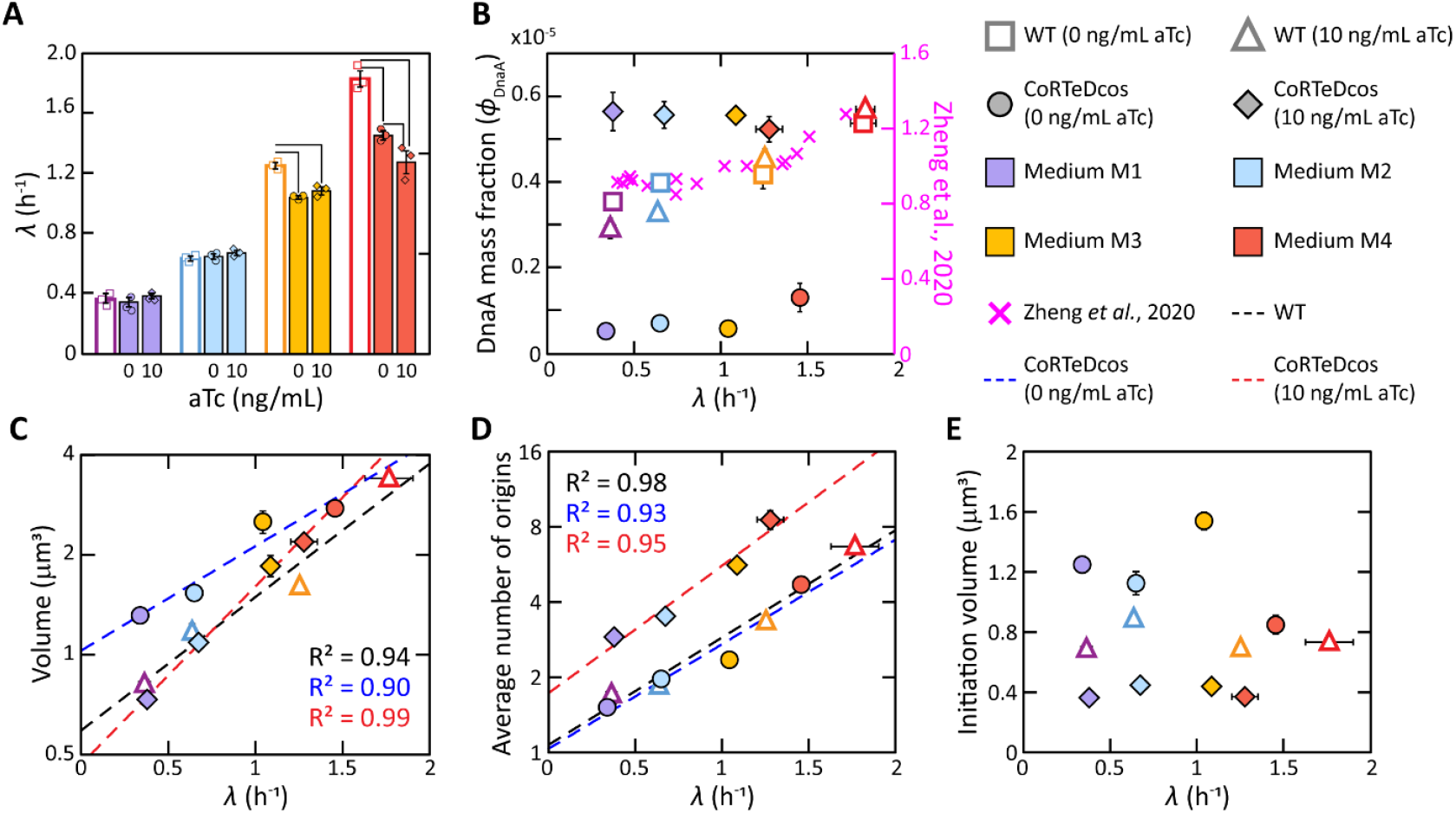
Phenotypical characterisation of *E. coli* CoRTeDcos. After balanced growth in turbidostat reactors, we characterised the **A)** growth rate, **B)** mass fraction of DnaA (*ϕ*_DnaA_), **C)** volume, **D)** number of origins and **E)** volume of initiation of *E. coli* MG1655 (open symbols) and *E. coli* CoRTeDcos (filled symbols) in four different media (M1: purple, M2: blue; M3: yellow; M4: red) and two concentrations of aTc (0 ng/mL and 10 ng/mL). Each bar or point is the average of three independent biological replicates (i.e., separate turbidostat cultivations), while the error bar indicates the standard deviation. In **A)**, individual data points are plotted and statistically significant differences (t-test, *p* < 0.05) are indicated by the black lines. In **B)**, data extracted from Zheng *et al*.^*12*^ is plotted for comparison. In **C)** and **D)**, cellular parameters are plotted on semi-log axes and exponential fits were applied to indicate relationships between growth rate and different parameters for either *E. coli* MG1655 (black line) or *E. coli* CoRTeDcos (blue line: 0 ng/mL; red line: 10 ng/mL).

*E. coli* CoRTeDcos recapitulates many known general relationships between growth rate and cell cycle even in the absence of DnaA-ATP/DnaA-ADP interconversion. Both cellular volume (Figure 2C) and number of origins (Figure 2D) scaled exponentially with growth rate, in agreement with the general growth law^7,12,42^. At the same time, the volume of initiation remained roughly constant (Figure 2E). The increase in *ϕ*_DnaAcos_ due to the presence of aTc had a clear effect on all these parameters. Specifically, larger *ϕ*_DnaAcos_ led to a decrease in volume and an increase in the number of origins. Even in this new condition, both parameters still increased exponentially with growth rate. The volume of initiation of *E. coli* CoRTeDcos cultured in 10 ng/mL of aTc was lower than the one obtained in absence of aTc and remained roughly constant across growth rates. The observed effect of an increase in *ϕ*_DnaAcos_ is consistent with the predicted behaviour of a previously described model of titration-based control of DNA replication^25^. In this type of systems, the initiation size is set by the accumulation of enough DnaA to saturate the high affinity motifs present on the chromosome^25,43^. Thus, increased synthesis of DnaAcos leads to faster occupation of these titration sites and earlier initiation events, reducing the average volume and increasing the number of origins of the population.

### *E. coli* CoRTeDcos DNA replication initiation behaves as a titration-based system

After observing growth-related phenotypes, we decided to further investigate the response of *E. coli* CoRTeDcos at varying aTc concentrations. We selected media M1, leading to slow growth, and M3, which yields a condition of fast growth with overlapping replication forks. We then used a total of eight different aTc concentrations to vary the *ϕ*_DnaAcos_ and probe its effect on the strain physiology.

As expected, elevated concentrations of aTc led to increases in the abundance of DnaAcos (Figure 3A). Furthermore, the obtained DnaAcos expression levels as function of aTc concentrations were comparable between media M1 and M3, further validating the capacity of the co-regulatory construct to maintain stable *ϕ*_DnaAcos_ across growth rates. The different concentrations of aTc all led to similar growth rates in the same medium (Figure S4A). The volume and number of origins of *E. coli* CoRTeDcos were considerably different at the same *ϕ*_DnaAcos_ across media (Figure 3B-C, see also Figure S4B-C for media M2 and M4), further reinforcing the idea of the general growth law that these parameters are set by the specific nutrient availability^7,12,42^. We also noticed an exponential decrease of cellular volume within the same medium at increasing *ϕ*_DnaAcos_ values (Figure 3B), while the average number of origins increased exponentially (Figure 3C). These tight correlations span a large range of DnaAcos protein levels, ultimately breaking down at the highest tested aTc concentrations. The anti-correlation between volume and number of origins was lost in medium M1 supplemented with 20 ng/mL of aTc, where both values increased. In medium M3, concentrations of 15 and 20 ng/mL caused an unexpected increase of cell volume and a decrease in the number of origins.

**Figure 3.**
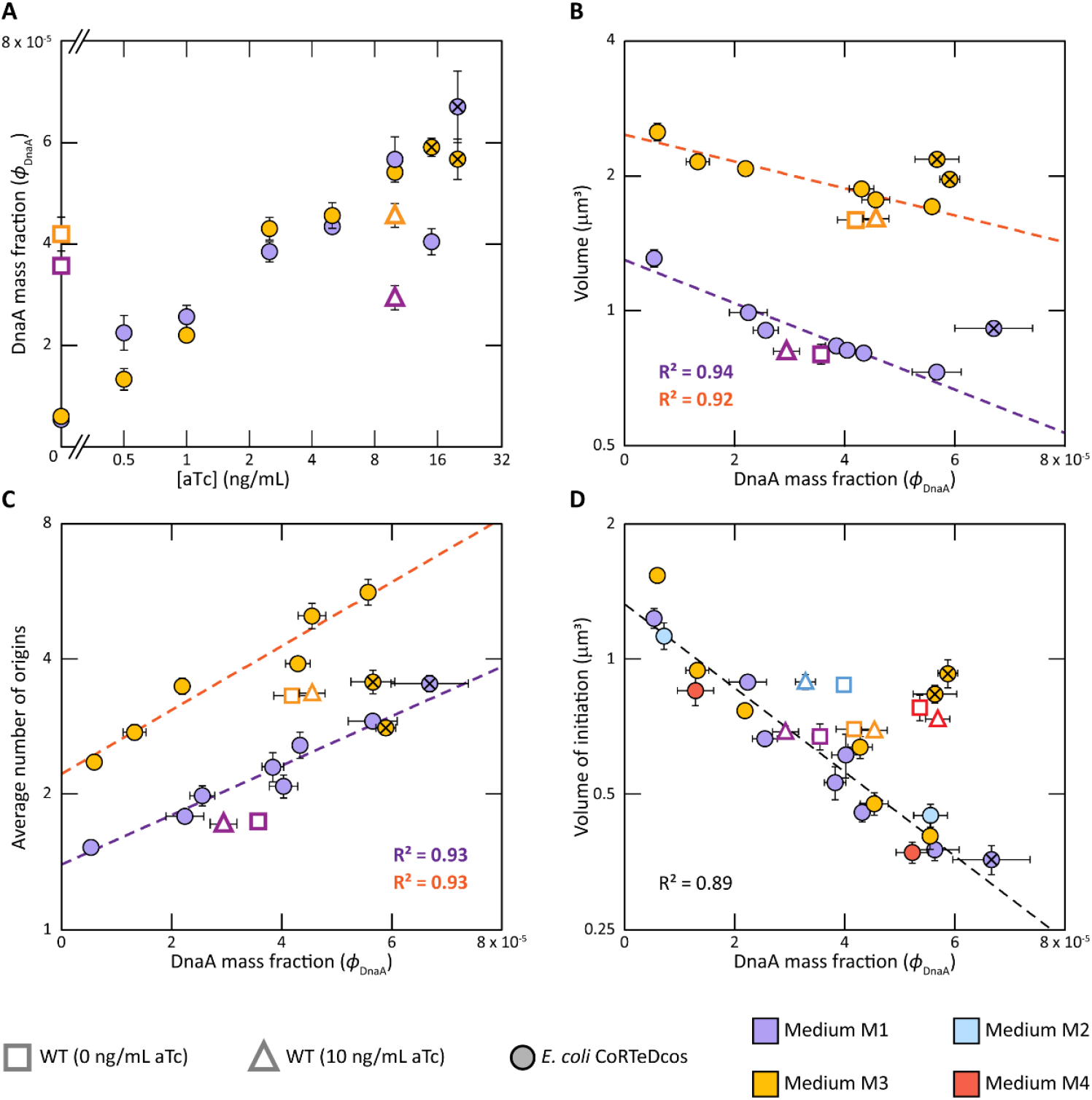
The influence of the abundance of DnaAcos on volume, number of origins and volume of initiation in *E. coli* CoRTeDcos. **A)** Values of *ϕ*_*DnaAcos*,_ at different concentrations of aTc. **B)** Cell volume, **C)** number of origins and **D)** volume of initiation per origin of *E. coli* CoRTeDcos (filled circles) are all correlated to *ϕ*_*DnaAcos*_. In **D)**, data coming from *E. coli* CoRTeDcos grown in media M2 and M4 is also presented. For other panels, data coming from media M2 and M4 is plotted in Supplementary Information (Figure S4D, S4E and S4F). In all graphs, *E. coli* MG1655 values are shown for comparison (open symbols). Purple, blue, yellow and red symbols represent data acquired from cells grown in media M1, M2, M3 and M4, respectively. The dashed lines are exponential fits of *E. coli* CoRTeDcos values. Crossed points represent the outliers excluded from the exponential fits. In **D)**, a single exponential trendline was used to fit data coming from all media. See Supplementary Information (Figure S4B) for individual trendlines for media M1 and M3. In all cases, cells grown in M1 with 20 ng/mL of aTc and in M3 with 15 and 20 ng/mL of aTc were deemed as outliers due to the discrepancy in the change in volume compared to the general trend. Cell volume plots where these conditions are also used are available in Supplementary Information (Figure S4C). All data points are the average of three independent biological replicates and error bars represent the standard deviation. In some cases, error bars are smaller than the symbols.

On the other hand, the volume of initiation seemed to be independent of growth rate and uniquely determined by the abundance of DnaAcos (Figure 3D, Figure S4D). More specifically, a single exponential relationship can explain the change in volume of initiation of *E. coli* CoRTeDcos caused by different *ϕ*_DnaAcos_ (Figure 3D, Figure S4B). The volumes of initiation of *E. coli* CoRTeDcos in media M3 at 15 and 20 ng/mL of aTc emerged as outliers.

Finally, we compared the values of volume of initiation of *E. coli* CoRTeDcos at different *ϕ*_DnaAcos_ with predictions coming from a titration model^25^ and a titration-switch model^25^. We also included predictions coming from the recently proposed titration-extrusion and titration-switch-extrusion models^44^, where the nucleoid-associated protein H-NS facilitates replication initiation by extruding DnaA from the chromosome. To this end, we simulated the expression, titration and/or interconversion of DnaA in the cell over time, together with the expression and effect of H-NS for models including extrusion^46^. When model-specific conditions causing DNA replication initiation are met (e.g., saturation of titration sites in a titration model), the volume of initiation was recorded. The synthesis rate of DnaA was then systematically changed. For both prediction and experiments, values of volume of initiation were normalised to the average of all values to obtain relative volumes of initiation. Relative DnaA expression levels were calculated in the same way. The simulations were performed for four different doubling times, similar to the ones experimentally obtained (i.e., 120 min, 60 min, 30 min and 20 min). The experimental datasets were then compared to the predictions of the different models at the same doubling time via reduced chi-squared statistics 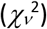 (see also Methods for more details on the simulations and statistical analysis). We noticed that the experimental data resembled predictions coming from a pure titration model more closely (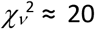, Figure 4) and deviated more considerably from predictions coming from a titration-switch model^25,44^ (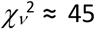, Figure S5A) or the recently proposed titration-extrusion^44^ (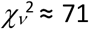, Figure S5B) and titration-switch-extrusion models^44^ (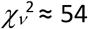, Figure S5C). The values previously mentioned as outliers (i.e., 20 ng/mL of aTc in medium M1 and 15 and 20 ng/mL of aTc in medium M3) were also included in the statistical analysis, but the results did not change when excluding them (see Source Data of Figure S5 for the related 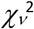 values). Notably, a recent study altered the mRNA levels of wild-type *dnaA* and found that the relationship between volume of initiation and DnaA expression does not support a model based solely on initiator titration^44^. We attribute the difference between this recent report and our study to the absence of DnaA interconversion in *E. coli* CoRTeDcos. We refrained from directly comparing the two datasets because changes in mRNA levels do not directly lead to the same changes at a protein level^45^.

**Figure 4.**
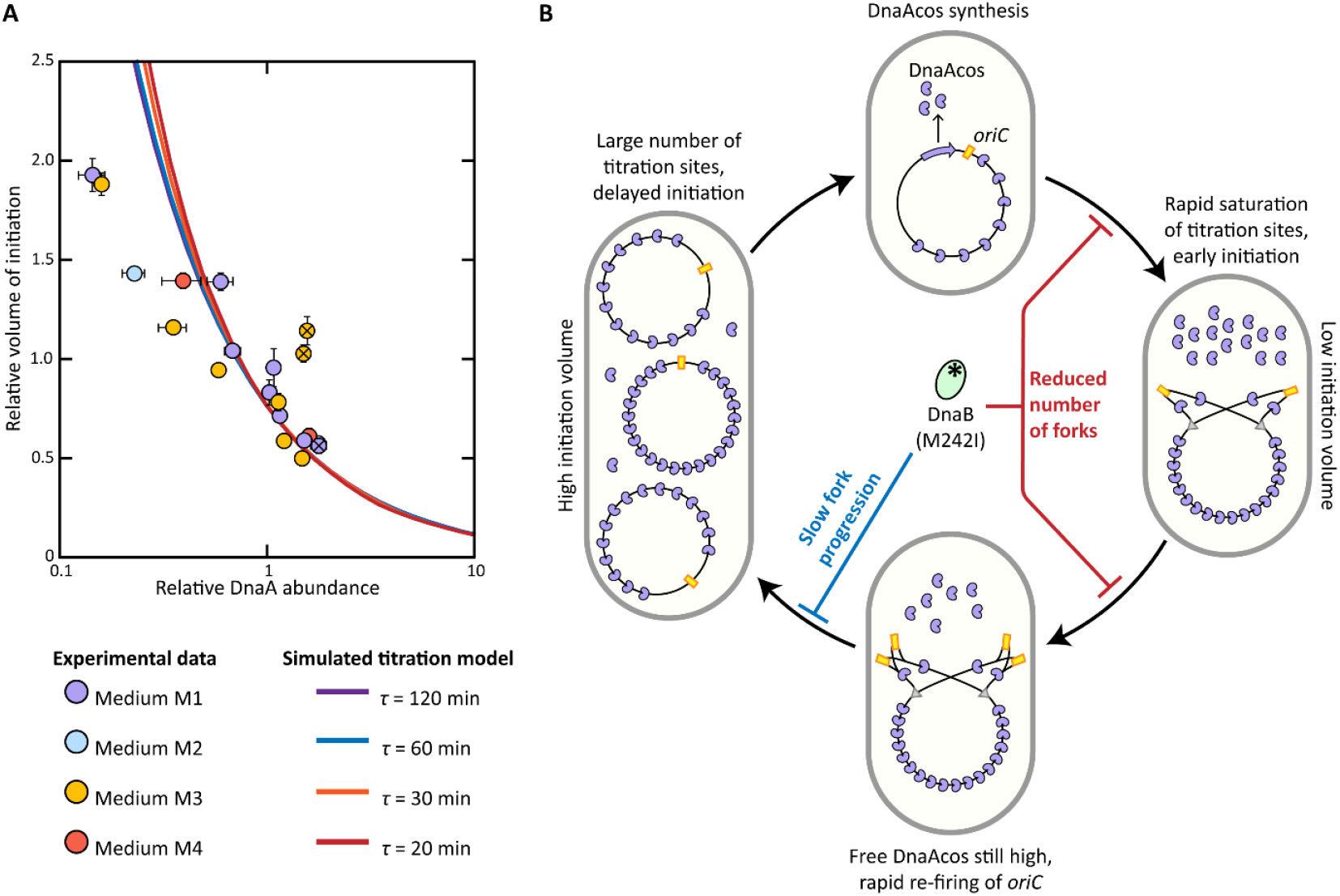
Behaviour and limitations of a titration-based control of DNA replication. **A)** Comparison of the relationship between relative change in DnaA expression and relative change in volume of initiation from *E. coli* CoRTeDcos measurements (circles) and predictions of a titration model (smooth lines). Simulations were performed using a doubling time of either 120 min (purple line), 60 min (blue line), 30 min (orange line) or 20 min (red line). Purple, blue, yellow and red symbols represent data acquired from cells grown in media M1, M2, M3 and M4, respectively. Crossed points represent the outliers previously mentioned in Figure 3. All data points are the average of three independent biological replicates and error bars represent the standard deviation. In some cases, error bars are smaller than the symbols. **B)** The predicted instability of titration-based systems at high growth rates and concentrations of DnaAcos leads to cells oscillating between a low initiation volume (right) and a high initiation volume (left). We hypothesise that the M242I mutation in DnaB rescues the cell from lethal hyper-replicative stress by limiting initiations (red lines) and/or lowering fork collisions (blue line).

Notably, there were two conditions in which the behaviour of *E. coli* CoRTeDcos differed from the trendline predicted by a titration system. First, cell populations grown in absence of aTc, and thus with low DnaA concentrations (Figure 2B, 3A), showed a lower volume of initiation than expected from simulations of the titration model (Figure 4A). This difference might hint at additional unknown regulatory mechanisms stabilising initiation. Furthermore, the absence of aTc was also associated with large-scale proteomic changes in *E. coli* CoRTeDcos compared to conditions where the inducer was present, hinting at a complex phenotype that cannot be easily explained via DnaAcos abundances alone (Figure S6). The second case of deviation from the predicted trend concerns the behaviour of *E. coli* CoRTeDcos in medium M3 with high concentrations of aTc. We explain this phenomenon with the predicted instability of titration-based systems at high growth rates^25,26^ (Figure 4B). In nutrient-rich conditions leading to overlapping replication forks, the rate of new DnaA synthesis is higher than the rate at which new DnaA boxes are formed per origin^24,25^. As a result, the available titration sites are quickly saturated and the free DnaA concentration increases rapidly, causing an early initiation event. At this stage, the free DnaA concentration remains high due to the nutrient-imposed synthesis rate and one or more new replication rounds are rapidly re-initiated. With the progression of multiple replication forks, the total synthesis of titration sites, summed over all origins, becomes higher than DnaA synthesis rate. The resulting accumulation of a large titration power leads to prolonged periods of low free DnaA concentrations and no new initiation events. Each cell cycle thus exhibits multiple rounds of replication initiation, causing the initiation volume per origin to oscillate between a high and a low value^25,26^. This phenomenon is normally associated with lethal hyper-replicative stress^4,28,30^ (Figure 1B). In *E. coli* CoRTeDcos, the pressure against this stress possibly caused the selection of a mutation in DnaB. We hypothesise that the lower processivity of the helicase resulting from the M242I mutation^37^ has a two-fold function. First, a slower progression on the chromosome might reduce the chance of two forks colliding when a replisome is stalled, minimising DNA damage. Second, the DnaB mutation can limit the total number of ongoing replication forks. DnaB is known to be present in few copies inside *E. coli* cells^46^ and six monomers are needed to form a DNA helicase^47^. For this reason, a slower processivity of the DNA helicase might lead to a longer turnover time before DnaB subunits become available to start a new round of replication, independently on whether DnaAcos already unwound *oriC*. The combination of these two effects might stabilise the system and allow *E. coli* CoRTeDcos to grow in all the tested conditions. However, cells might still be exhibiting oscillating replication initiations and the type of measurements employed here would only reflect population averages. Our observations in medium M3 at the highest concentrations of inducer (i.e., higher volumes (Fig. 3B) and lower number of origins (Fig. 3C)) could then indicate that cells spend on average more time in the phase in which no replicative events take place.

### *E. coli* CoRTeDcos proteome suggests hyper-replicative stress during fast growth

To obtain further evidence on the instability of titration-based systems during fast growth, we looked for changes in the proteome of *E. coli* CoRTeDcos indicative of a condition of hyper-replicative stress. Although there is no characterised proteome signature for this type of stress, the events eventually leading to cell death in these conditions are well known (Figure 5A). Uncontrolled initiations of replication lead to several parallel forks accumulating on the chromosome. The quick depletion of the dNTP pool in the cell then causes the replication forks to slow down and eventually arrest and/or collide^48,49^. Moreover, the presence of reactive oxygen species (ROS) in the cell leads to the introduction of lesions in the nascent DNA^50^. For these reasons, mutations lowering origin firing^35,51^, increasing the available amount of dNTPs^49^ or decreasing the presence of ROS^36,50^ are known to rescue the cell from hyper-replicative stress. As such, we investigated the proteome of *E. coli* CoRTeDcos for changes in the expression of proteins that would reduce initiation events (DiaA^34^, DnaB^34^, SeqA^35^, YfdQ^51^), increase the dNTP pool (Nrd proteins)^35,48,49^ or reduce the formation of ROS (AtpAB, CydAB, Fre, IscU)^36^ (Figure 5B). We also searched for changes in the synthesis of proteins associated with DNA repair mechanisms, as we argued that they could reveal the presence of DNA damage.

**Figure 5.**
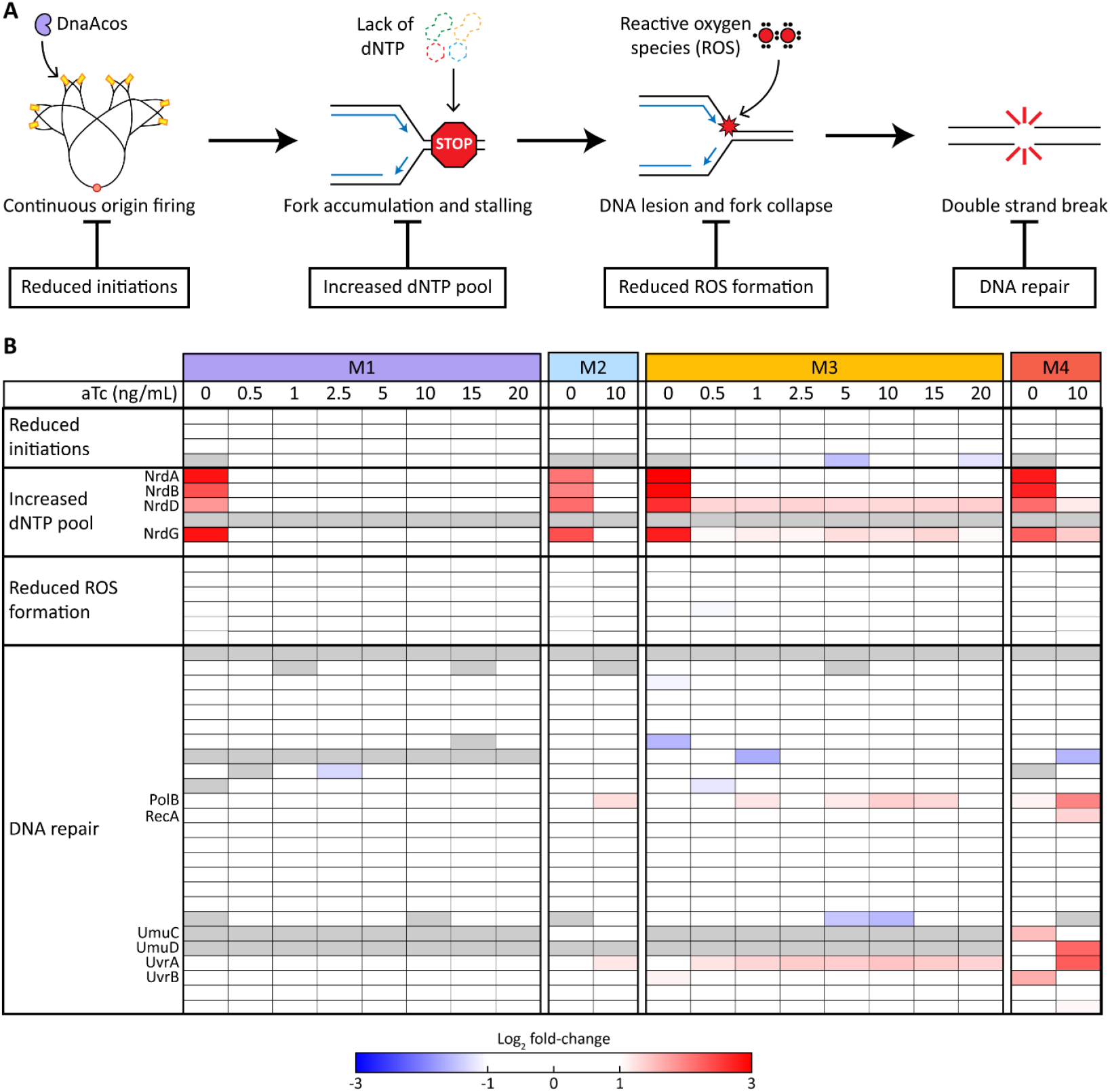
*E. coli* CoRTeDcos undergoes replicative stress during fast growth. **A)** Hyper-replicative stress caused by continuous origin firing leads to fork accumulation and stalling, ROS-mediated DNA lesions and fork collapse. These events ultimately generate double strands breaks in the chromosome of *E. coli* that, if not repaired, lead to cell death. Several proteins can suppress this lethal phenotype, by either reducing the number of initiations, increasing the dNTP pool, reducing the formation of ROS or repairing DNA. **B)** Changes in production of sets of proteins involved in the different forms of rescue from hyper-replicative stress. For each cell in the heatmap, proteins produced with more than a two-fold difference compared to wild-type are mapped in gradients of either blue (under-expression) or red (over-expression). Proteins that could not be detected in more than two samples either in *E. coli* CoRTeDcos or in *E. coli* MG1655 are indicated in grey and were omitted from the analysis. Proteins with a significantly different production compared to the wild-type background are indicated in the figure. The complete list of proteins belonging to each group is available in the Source Data of the study, together with their relative fold-change.

In the slowest tested growth conditions (medium M1, *λ* ≈ 0.37 h^-1^), we noticed no proteome changes that could indicate an adaptation to replicative stress. With richer nutrient conditions and thus improved growth rates, we observed a progressively larger overexpression of proteins involved in dNTP synthesis and DNA repair. In medium M2 (*λ* ≈ 0.7 h^-1^), the presence of aTc led to an increased abundance of two DNA repair proteins (PolB^52,53^, UvrA^54^). In media promoting overlapping replication forks (i.e., M3 and M4), two additional proteins involved in DNA repair (UmuC^55,56^, UvrB^54^) were overproduced even in absence of aTc. Increased levels of inducer and thus DnaAcos then further aggravated these proteome changes. In general, we observed a more abundant production of proteins involved in repairing DNA lesions and double-strand breaks (PolB^52,53^, RecA^57^, UvrABD^54^, UmuCD^55,56^). The anaerobic ribonucleotide reductase protein NrdD and its activator protein NrdG were also among the proteins overexpressed upon induction of DnaAcos, despite the fact that all our cultivations were performed in aerobic conditions. A similar response has been previously reported in hyper-replicating mutants^34,49^. The NrdA, NrdB, NrdD and NrdG proteins were also strongly overexpressed (between 3.6 and 8-fold) in absence of aTc at all growth rates. The *nrdA* and *nrdD* operons are known to be part of the DnaA regulon^58,59^, yet their activation or repression only partially depends on DnaA^58,60^. It is thus possible that very low *ϕ*_DnaAcos_ (between 0.13 and 0.23 times the wild type levels) contribute to elevated transcription of the *nrd* operons.

We did not notice any major changes in expression that could indicate a reduction in DNA replication initiation or a lower formation of ROS. Moreover, the observed changes were not attributable to the presence of aTc alone, as evident from the analysis of wild-type *E. coli* proteomes (Figure S6). Overall, the observed changes in the proteome of *E. coli* CoRTeDcos seem to indicate the presence of DNA damage during faster growth at almost all inducer concentrations. Moreover, they suggest that this type of stress can be present even in cells that do not lose the typical anti-correlation between cell volume and the number of origins.

## Discussion

In this study, we established a titration-based control of DNA replication in *E. coli* CoRTeDcos with only minimal genetic modifications and the aid of selective pressure. This result was achieved (i) by replacing the ATP/ADP-controlled wild-type DnaA initiator with an ‘always-active’ variant DnaAcos, and (ii) by selection of a single aminoacidic substitution in the helicase protein DnaB. The use of a co-regulatory construct further allowed us to probe the behaviour and limits of the system at different concentrations of DnaAcos (Figure 2 and 3). Titration of DnaAcos relied entirely on the native amount of DnaA boxes, further confirming the capacity of *E. coli* chromosome to sequester the initiator protein^24^. Most importantly, we obtained a titration-based system recreating the phenomenon of stable volumes of initiation across growth rates (Figure 3 and 4). In *E. coli* CoRTeDcos, the specific value of volume of initiation can then be controlled by inducing the *P*_*LtetO-1*_ promoter via aTc.

On top of providing insights into the capacity and limit of DnaA titration in replication control, it is interesting to look at our results from an evolutionary perspective. Initiator titration has been proposed as the possible control mechanism of slow-growing ancestors of *E. coli*^26,61^. In this hypothesis, the interconversion between DnaA-ATP/DnaA-ADP was adopted later in this bacterium evolutionary trajectory to allow doubling times shorter than the time needed for chromosome replication. In line with this idea, we previously reported a shared genomic configuration favouring the titration of DnaA among most *E. coli* strains^24^. Here, we elaborate on previous studies by showing that our *E. coli* CoRTeDcos strain seems to be more stable during slow growth. In this condition, the volume of initiation maintained an exponential inverse relationship with *ϕ*_DnaAcos_ (Figure 3) and we did not observe any change in protein expression hinting at the presence of DNA damage (Figure 5). On the other hand, growth conditions promoting overlapping replication forks caused atypical behaviours that can be reconciled with hyper-replicative stress. Several proteins involved in nucleotide synthesis and DNA repair pathways were always overproduced (Figure 5) and high levels of DnaAcos caused a counterintuitive increase in volume of initiation (Figure 4). We speculate that this same instability served as evolutionary pressure leading to the adoption of the DnaA interconversion mechanism in *E. coli* ancestors. It is important to note that our conclusions are based on population averages that inevitably mask any single-cell behaviour or variation. For this reason, we cannot directly observe the occurrence of multiple initiation events within the same cell division cycle predicted by the titration model during regimes of overlapping replication. Notably, unstable regimes could also emerge during slow growth in absence of overlapping replication forks and at high *ϕ*_DnaAcos_, but remain undetectable by our approach. In the future, studying *E. coli* CoRTeDcos in microfluidic cultivation settings^6,62^ could shed light on the intricacies of its regulation strategy at a single-cell level and further help in tuning the system.

Finally, we believe that our results may be useful in the field of synthetic cells. Specifically, the creation of a functional synthetic cell must meet the fundamental requirement of one replication event per cell division^63^. If not, the cellular DNA content would not remain constant throughout generations, continually increasing until death^63^. This requirement is already not trivial in natural cells^64^, but its recreation faces further technical limitations in the field of synthetic cells. Due to the complexity of building synthetic cells with a bottom-up approach, the use of minimal systems is generally preferred in the reconstitution of cellular process^63,65^. Hence, simple replication machineries and control strategies are preferred for a bottom-up synthetic cells. The design criteria employed here can in principle be repurposed for initiator proteins different from DnaA, to allow a simple control strategy for other, more minimal replication machineries. For instance, the replicative machinery derived from bacteriophage *ϕ*29^66^ has been the system of choice in the study of DNA replication for synthetic cells in the last years^63,67^. Engineering high- and low-affinity binding sites for the terminal DNA-binding protein (p3) would thus yield a titration-based control mechanism that only requires addition of binding motifs in specific sections of the future synthetic cell chromosome^68^. The resulting system would allow streamlined and growth-coupled DNA replication, without the input of additional ATP^13^ or eukaryotic-like complex genetic architectures^69^.

## Materials and methods

### Bacterial strains and growth conditions

A list of bacterial strains used in this study is provided in Supplementary Information (Table S1). *Escherichia coli* DH5α (NEB) was used for general plasmid propagation and standard molecular techniques. The wild-type *E. coli* MG1655 and the mutant strain *E. coli* CoRTeDcos were used for all phenotypical characterisation experiments.

*E. coli* CoRTeDcos was routinely propagated at 30 °C or 37 °C and 200 rpm in M9 liquid medium supplemented with either 22 mM of glycerol or 22 mM glucose. All other *E. coli* strains were routinely propagated at 30 °C or 37 °C and 200 rpm in LB liquid medium. For constant density cultivations, four different liquid media were used (M1 to M4), yielding growth rates ranging from 0.4 h^-1^ to 1.8 h^-1^. Media M1, M2 and M3 were based on M9 medium, supplemented with either glycerol or glucose as carbon source and eventually different amounts of amino acids. Medium M4 was a tryptic soy broth. Complete medium composition is available in Supplementary Information (Table S2). All strains were cultivated at 37 °C in either M1, M2, M3 or M4 media, eventually supplemented with anhydrotetracycline (aTc). Media were eventually supplemented with agar (15 g/L) to obtain solid media and with antibiotics for strain selection or plasmid propagation (50 mg/L kanamycin, 100 mg/L ampicillin, 25 mg/L chloramphenicol).

Phosphate-buffered saline solution (PBS) (137 mM NaCl, 2.7 mM KCl, 10 mM Na_2_HPO_4_, 1.8 mM KH_2_PO_4_) was used to wash cells during sample preparation.

### Plasmid and linear knock-in fragment construction

A list of all plasmids used in this study is provided in Supplementary Information (Table S3). The CoRTeDcos mutation was introduced in *E. coli* MG1655 using the pSC020 plasmid system, containing the *λ*-red recombineering system and the Cre recombinase^70^. To obtain the linear fragment for recombination while avoiding toxicity of DnaAcos, the CoRTeDcos locus was split into two fragments, each cloned in a separate plasmid. The pCoRe-T plasmid was based on a pUC19 backbone and contained, in order, a chloramphenicol resistance cassette flanked by *lox66* and *lox71* sites^71^, the TetR repressor under the control of the *P*_*LtetO-1*_ promoter and a BsaI restriction site generating a 5’-TCAG sticky end. The pCoRe-Dcos plasmid was based on a pUC19 backbone and contained, in order, a BsaI site generating a 5’-AGTC sticky end, the *dnaAcos* gene and a kanamycin resistance cassette flanked by *lox66* and *lox71* sites^71^. The DnaAcos gene was chemically synthesised (BG22166). To obtain the linear recombination fragment, first a PCR reaction was performed on pCoRe-T with primers BG22660 and BG20764 and on pCoRe-Dcos with primers BG21406 and BG22661. The two fragments were then purified and mixed in equimolar amounts together with BsaI (NEB) restriction enzyme and T4 DNA ligase (NEB) in CutSmart reaction buffer (NEB). The reaction mixture was incubated at 37 °C for 1 hour, before being heated up to 80 °C for 5 min to inactivate the enzymes. 5 μL of reaction were then used as PCR template with primers BG22660 and BG22661 to amplify the full recombination fragment generated in the previous restriction-ligation reaction. Primer BG22660 contained an overhang with 50 bp of homology to positions 3883943-3883993 of *E. coli* MG1655 genome (upstream of the *P*_*dnaA*_ promoter); primer BG22661 contained an overhang with 50 bp of homology to positions 3882292-3882342 of *E. coli* MG1655 genome (downstream of *dnaA* and into the *dnaN* gene).

DNA fragments were amplified by PCR using Q5® High-Fidelity DNA Polymerase (NEB) following manufacturer instructions. Specific oligonucleotides and synthetic DNA were designed and synthesised (IDT) to introduce proper overhangs for assembly. The sequences of primers and synthetic DNA used in this study are provided in Supplementary Information (Table S4). Plasmid assembly was performed using NEBuilder® HiFi DNA Assembly Master Mix (NEB).

### Transformation procedure and creation of *E. coli* CoRTeDcos

For all transformation procedures, cells were made electrocompetent and transformed through electroporation as previously described^24^. The pCoRe-T and pCoRe-Dcos plasmids were transformed and propagated in *E. coli* DH5α. The pSC020 plasmid^72^, containing the *λ*-red recombination system and the Cre recombinase, was transformed in *E. coli* MG1655. To create *E. coli* CoRTeDcos, electrocompetent *E. coli* MG1655 cells carrying the pSC020 plasmid were transformed with 200 ng of linear CoRTeDcos fragment, plated in solid M1 medium supplemented with kanamycin, chloramphenicol and ampicillin, and incubated at 30 °C. Insertion of the co-regulatory construct was confirmed first by colony PCR using primers BG22467 and BG22496 and then by Sanger sequencing of the locus. Correct colonies were propagated in solid M1 medium supplemented with ampicillin and 1 mM IPTG to induce expression of Cre recombinase, and incubated at 30 °C. Colonies that lost kanamycin and chloramphenicol resistances were propagated in solid M1 medium at 37 °C to promote loss of the pSC020 plasmid. After screening for colonies sensitive to ampicillin, whole genome sequencing was performed to ensure that the co-regulatory construct was the only source of DnaA in the chromosome of *E. coli*. The final strain *E. coli* CoRTeDcos emerged during prolonged incubation of the initial mutant strain in M3 medium supplemented with 10 ng/mL of aTc. Whole genome sequencing was performed to identify the *dnaB* mutation selected for in *E. coli* CoRTeDcos (Table S5).

### Turbidostat cultivation of cell cultures and growth characterisation

We employed constant-density cultivation inside turbidostat reactors to obtain *E. coli* cultures in balanced growth. *E. coli* strains were first grown overnight in 5 mL of M1 or M3 medium at 37 °C and 180 rpm. The densely-grown culture was then used to start a turbidostat cultivation using the Pioreactor platform (Pioreactor 40 mL v1.0; software version 25.5.22; Pioreactor inc., https://pioreactor.com). Experiments were started according to manufacturer instructions for fluidic lines, electrical connections and user interface. Cultures were started at an OD_600nm_ of 0.05 in media M1-M4, supplemented with aTc appropriately. Default settings were employed for frequency of optical density measurements; stirring was set to 700 rpm. The target optical density was set to OD_600nm_ 0.3 and the media reservoir was filled with the same medium as in the cultivation chamber. Cultivation medium was pumped in the vessel from the reservoir to maintain a constant optical density. Cells were balanced in exponential growth at the target optical density for at least ten generations before harvesting. Finally, cells were harvested in different amounts for downstream processing. Three biological replicates were obtained for every strain and condition by performing independent turbidostat cultivations.

For each turbidostat cultivation, we estimated growth rate (*λ*) using the dilution rate (*D*). The total volume (*V*_SteadyState_) and time (*t*_SteadyState_) spent at the steady-state optical density were used to calculate the flow rate 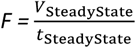. *F* was then used together with the cultivation volume (*V*_Culture_= 20 mL) to calculate dilution rate and thus growth rate, following 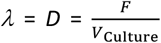. Doubling times (*τ*) were obtained from growth rates measurements.

In microplate reader growth assays, cells coming from an overnight culture were used to inoculate a medium containing different aTc concentrations at an initial OD_600nm_ of 0.05. 150 µL of inoculated medium was topped with 50 µL of mineral oil to prevent evaporation during the growth experiment. Growth of the different strains was assessed in each condition in three biological replicates. The medium used for the overnight culture was the same medium used in the final microplate reader assay, with the difference that cells were always propagated in absence of aTc overnight. The two media used were both based on M9, either supplemented with 20 mM of acetate (slow growth) or with 20 mM of glucose and RPMI amino acids (fast growth). Growth was then monitored by absorbance at 600 nm using a Synergy H1 microplate reader (BioTek×) either over 48 hours for slow growth or 20 hours for fast growth experiments. A previously published MATLAB script^73^ was used to estimate growth rate from each growth curve.

### Proteomic analysis

To perform proteomics, 10 mL of turbidostat cultures of *E. coli* strains in different growth conditions were harvested. Cells were then washed in PBS before being pelleted and bacteria were lysed with 100% trifluoracetic acid (TFA) according to the SPEED protocol^74^ with slight adaptations^75^. Briefly, 25 μL of 100% TFA was added to the pellet, incubated for 5-7 min at 55 °C and neutralised with 225 μL of 2 M Tris. Protein concentration from lysed samples were quantified via Bradford assay. 50 μg total protein amount per sample were reduced and alkylated (9 mM tris(2-carboxyethyl)phosphine (TCEP) and 40 mM chloroacetamide (CAA)). Protein digestion was performed with a Trypsin (Roche)-to-protein ratio of 1:50 overnight at 37 °C. Samples were desalted by solid-phase extraction (C18) and stored after lyophilisation at -80 °C. Peptides were analysed on a Vanquish Neo UHPLC (micro-flow configuration; Thermo Fisher Scientific) coupled to an Orbitrap Exploris 480 mass spectrometer (Thermo Fisher Scientific). Around 25 μg peptides were applied directly onto a commercially available Acclaim PepMap 100 C18 column (2 μm particle size, 1 mm ID x 150 mm, 100 Å pore size; Thermo Fisher Scientific) and separated using a 60 min linear gradient ranging from 3% to 28% solvent B (0.1% FA, 3% DMSO in ACN) in solvent A (0.1% FA, 3% DMSO in HPLC grade water) at a flow rate of 50 μL/min. The mass spectrometer was operated in data-independent acquisition (DIA) mode. MS1 full scans (360 – 1300 m/z) were acquired with a resolution of 120,000, an AGC target value of 1E6 and a maximum injection time of 50 ms. The cycle time was set to 3 s. MS2 scans (200 – 1800 m/z) were acquired over 40 DIA segments with widths adjusted to the precursor density (see Supplementary Table 8 in Wu *et al*.^76^). The scan resolution in the Orbitrap was set to 15,000 with an AGC target value of 1E5 and a maximum injection time of 22 ms. The HCD collision energy was set to 30%. The mass spectrometric raw files were analyzed with DIA-NN^77^ (version 1.8) using the Uniprot reference fasta file for *E. coli* K-12 (UP000000625, download 04.06.2023, 4403 protein entries) appended with common contaminants. FASTA digest for library-free search and deep learning-based spectra was enabled. N-term M excision, C carbamidomethylation and Ox(M) were activated. The peptide length range was set from 7 to 30, precursor charge ranged from 2 to 6, precursor m/z ranged from 360 to 1300 and fragment ion m/z ranged from 200 to 1800. Usage of isotopologues and match-between runs (MBR) were enabled. Precursor FDR was set to 1%. All other parameters were kept as default. The DIA-NN main report table “report.pg_matrix.tsv” was processed with the R-script DIAgui^78^, to obtain intensity-based protein quantification values (iBAQ intensities)^79^ for the determination of protein mass fractions, as well as label-free quantification values (MaxLFQ intensities)^80^ for relative protein quantification. Further processing was carried out with the in-house proteomics analysis tool “baybiomsCruncher” (https://github.com/nava20ir/baybiomsCruncher)^81^. This processing included filtering proteins for being quantified in all three replicate measurements of at least one experimental condition, as well as imputation of missing values by replacement with random values drawn from a normal down-shifted distribution (width = 0.3, downshift = 1.8). Protein mass fractions of DnaA variants in the proteome of *E. coli* MG1655 and *E. coli* CoRTeDcos were calculated by dividing the iBAQ value of DnaA for the summed iBAQ values of all detected proteins in a given sample. To estimate the change in the level of DnaA variants at different growth rates, we normalised the DnaA mass fraction obtained for each sample using the average DnaA fraction of the wild-type across growth rates. Ribosome fractions (*ϕ*_R_) were obtained by dividing the sum of the iBAQ value of all proteins associated with ribosomes for the summed iBAQ values of all detected proteins inside a given sample. To gain insights into the presence of hyper-replicative stress, MaxLFQ intensities of relevant proteins in *E. coli* CoRTeDcos were compared to the wild-type intensities. Only proteins that could be detected in at least two of the three biological replicates in both *E. coli* CoRTeDcos and *E. coli* MG1655 samples were considered. Protein fold-changes were used to generate a heat-map with a custom-made MATLAB script.

### Cell size measurements

Cell size measurements were estimated via phase-contrast microscopy. 3 mL of turbidostat cultures of *E. coli* were washed once with PBS and then resuspended in 10 μL of PBS in an Eppendorf tube. 1-2 μL of the final cell suspensions were immobilised on agarose pads (PBS supplemented with 1.5% agarose) on microscopy glass slides. Phase contrast images were acquired with an ECLIPSE Ti2 microscope (Nikon) equipped with a 100x immersion oil objective (N.A. = 1.4). Cells in the microscopic images were segmented using Omnipose^82^, yielding cell masks. A custom-written Python script was used to extract values for length and width in pixels of each identified cell from the cell masks. The camera pixel size (49.4 x 49.4 nm) was used to generate the corresponding measurements in µm. The rod shape of *E. coli* was approximated to a cylinder with a hemisphere at each pole and the volume was estimated as 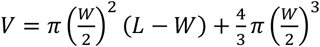, using the obtained length (*L*) and width (*W*). Generally, thousands of cells per replicate were used; exact numbers are reported in Supplementary Information (Table S6).

### Characterisation of C period

In *E. coli* cultures undergoing steady-state growth, the relative frequency of two genomic loci depends on the C period, their relative distance and the doubling time^7,12,83^. To estimate the length of C period, digital PCR was used to estimate the relative abundance of origins and termini of replication. To this end, 5 mL of turbidostat cultures of *E. coli* were washed with PBS before being pelleted. Genomic DNA was extracted using the DNeasy Blood & Tissue Kit (QIAGEN) and eluted in milliQ water. Genomic DNA concentration was assessed using Qubit dsDNA BR Assay Kit (Invitrogen). All samples were then diluted to a concentration of 5 ng/µL, before being diluted an additional 200 times. 2 µL of the resulting genomic DNA solution was then used as input in digital PCR reactions. Each reaction was performed with QIAcuity EvaGreen Mastermix (QIAGEN) in a 12 µL reaction, using a QIAcuity Digital PCR System (QIAGEN). Previously reported^7^ primer pairs were used to measure abundance of *oriC* (primers SJO416 and SJO417) and *ter* (primers SJO432 and SJO433).

The C period was then obtained as 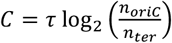, where 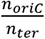 refers to the relative abundance of the origin of replication and the terminus as obtained through digital PCR, and *τ* refers to the culture doubling time, obtained from the growth rates from turbidostat cultivation.

### Flow cytometry estimation of genome equivalents

To perform flow cytometry, 2 mL of turbidostat cultures of *E. coli* were first added to 18 mL of ice-cold 70% V/V ethanol and allowed to fix at 4 °C for at least 12 h and for a maximum of 14 days^84^. Cells were washed once with PBS and then resuspended in 1 mL of PBS in an Eppendorf tube. 10 μL of the resulting cell suspension were stained with 2 μL of 100× Quant-iT× PicoGreen× dsDNA reagent (Invitrogen) in 25% DMSO (ratio 1:1). The cells were incubated together with the nucleic acid dye for at least 90 min at room temperature in the dark. Finally, 200 μL of PBS was added to the samples. All flow cytometry analyses were performed with a CytoFLEX (S) (Beckman Coulter Life Sciences) and the CytExpert software. For each sample, forward and side scatter measurements were obtained, together with emission filtered with a 530/30 nm transmission filter for DNA content measurement (B525-H channel, gain= 40). PicoGreen-stained particles were deemed to be bacterial cells and at least 100’000 of these events were collected per sample. The DNA content of cell populations was represented as a histogram versus fluorescence on the green channel. *E. coli* MG1655 cells grown in M1 and subjected to run-out replication (30 mg/mL cephalexin, 300 mg/mL cephalexin) were used to quantify the fluorescence intensity of one copy of *E. coli* chromosome as previously described^24,84^. The value obtained in this way was used to convert fluorescence intensity of experimental samples to genome equivalents. An example of the employed gating strategy is available in Supplementary Information (Figure S7).

### Characterisation of number of origins and volume of initiation

Theoretical analysis^7^ revealed a relationship between the length of the D period, the average genome equivalents 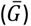, the length of the C period and the doubling time. Using this relationship and measured cellular parameters, the time of the cell cycle was obtained as 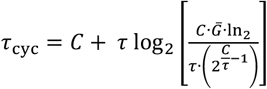. From here, the number of origins of the population (*noriC*) was estimated as 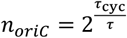. The volume of initiation (*V*^∗^) was estimated as 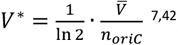, where 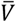 is the average cellular volume of the population.

### Simulations of titration, switch and extrusion models

To assess what theoretical model best fits our observations, we compared our experimental data with simulations of DNA replication in *E. coli* cells. To this end, we used a set of previously generated MATLAB scripts^44^ (https://github.com/BaiYangBqdq/dynamics_of_biomass_DNA_coordination/tree/main), based on theoretical works performed by other studies^23,25^ or by the same authors^44^. Briefly, the scripts simulate the expression, titration and/or interconversion of DnaA in the cell over time, together with the expression and effect of H-NS for models including extrusion. At every time step, the cell volume grows according to growth rate, the number and state of DnaA proteins (bound vs free; ATP-bound vs ADP-bound) are updated and model-specific parameters are compared to critical values that would trigger replication initiation. Every time the critical condition is reached, an initiation event is recorded and the relative initiation mass is saved. The simulation was repeated for a range of DnaA expressions levels by varying the DnaA synthesis rate and the cell doubling time. In this way, we generated predictions on the change in volume of initiation across different DnaA expression levels and growth conditions using either a titration model, a titration-switch model, a titration-extrusion model or a titration-switch-extrusion model. Due to the inability of DnaAcos to bind nucleotides^28–30^, we did not compare the experimental data with simulations of a switch-only model. We used the same value of the original work for all the parameters (see Table S7 in Supplementary Information for complete list), with the exception of the doubling time. The doubling time value was set to either 20 min, 30 min, 60 min or 120 min to approximate our experimental conditions. From our proteomics data, we observed that levels of the nucleoid-associated protein H-NS were roughly constant across the tested conditions. Therefore, we did not change the H-NS synthesis rate parameter when varying the cell doubling time. We refer to the original works for in depth mathematical derivations of the biological models and explanation of the scripts^23,25,44^. For both experimental and simulated data, the values of volume of initiation were normalised to the average of all values obtained in the same condition (i.e., the same medium for experimental data and same doubling time parameter for simulations) to obtain the relative volume of initiation. The relative DnaA values were obtained in the same way, starting from either *ϕ*_DnaAcos_ for experimental data or DnaA expression rate for simulations. The values of relative volume of initiation and relative DnaA obtained from the simulation were fitted to a trendline (*R*^2^ > 0.98) and the obtained equations were used to estimate values of relative volume of initiation at different values of relative DnaA. Reduced chi-squared statistics 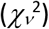 was then employed to identify how closely the experimental values resembled the predictions of the different models. The reduced chi-squared statistics was calculated as: 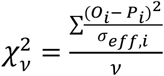, where *O* and *P* are experimental and predicted values, respectively, at the same relative DnaA value, *σ*_*eff,i*_is the expected variance and *ν* are the degrees of freedom. The expected variance was calculated as 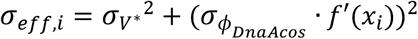, where *σ* ∗ is the standard deviation of the volume of initiation, 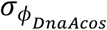 is the standard deviation of the relative *ϕ*_DnaAcos_ and *f*′(*x*_*i*_) is the slope of the trendline used to fit the predicted values at that value of relative *ϕ*_DnaAcos_. The degrees of freedom were calculated as *ν* = *N* - *k*, where *N* is the total number of experimental values and *k* is the total number of parameters of the equations used to fit the predicted values across doubling times. The model leading to the lowest 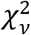 was considered as the one best describing the experimental observations. All experimental points were used in estimating the lowest 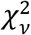 value, including the ones identified as outliers during the analysis of the behaviour of *E. coli* CoRTeDcos in media M1 and M3 and different aTc concentrations. The statistical analysis was also repeated excluding the outliers.

## Acknowledgements

We would like to thank Franziska Hackbarth and Tayma Midari for technical assistance and maintenance of mass spectrometers. We would like to thank Hans Heilig for technical assistance on performing all digital PCR experiments. We would like to thank Oscar Broström and David Fange for constructive discussions and a critical reading of the manuscript.

## Author contributions

Conceptualisation: LO, MB. Methodology: LO, DF, BAP. Software: DF, CL, JH. Investigation: LO, BAP. Formal analysis: LO. Visualisation: LO, BAP. Supervision: LO. Writing – original manuscript: LO, BAP. Writing – review & editing: BAP, MB, DF, CL, JH, RHJS, PRtW, JvdO, NJC, LO.

## Data and material availability

All data used in the study to generate plots, together with the raw proteomics data are available upon request by contact with the corresponding author.

## References

1. Ohbayashi, R. et al. Evolutionary Changes in DnaA-Dependent Chromosomal Replication in Cyanobacteria. Front Microbiol 11, 786 (2020).

2. Erzberger, J. P., Mott, M. L. & Berger, J. M. Structural basis for ATP-dependent DnaA assembly and replication-origin remodeling. Nat Struct Mol Biol 13, 676–683 (2006).

3. Leonard, A. C., Rao, P., Kadam, R. P. & Grimwade, J. E. Changing Perspectives on the Role of DnaA-ATP in Orisome Function and Timing Regulation. Front Microbiol 10, 2009 (2019).

4. Simmons, L. A., Breier, A. M., Cozzarelli, N. R. & Kaguni, J. M. Hyperinitiation of DNA replication in Escherichia coli leads to replication fork collapse and inviability. Molecular Microbiology 51, 349–358 (2004).

5. Fernandez-Fernandez, C., Gonzalez, D. & Collier, J. Regulation of the Activity of the Dual-Function DnaA Protein in Caulobacter crescentus. 6, e26028 (2011).

6. Wallden, M., Fange, D., Lundius, E. G., Baltekin, Ö. & Elf, J. The Synchronization of Replication and Division Cycles in Individual E. coli Cells. Cell 166, 729–739 (2016).

7. Si, F. et al. Invariance of Initiation Mass and Predictability of Cell Size in Escherichia coli. Current Biology 27, 1278–1287 (2017).

8. Donachie, W. D. Relationship between Cell Size and Time of Initiation of DNA Replication. Nature 1968 219:5158 219, 1077–1079 (1968).

9. Cooper, S. & Helmstetter, C. E. Chromosome replication and the division cycle of Escherichia coli Br. Journal of Molecular Biology 31, 519–540 (1968).

10. Helmstetter, C. E. & Cooper, S. DNA synthesis during the division cycle of rapidly growing Escherichia coli B/r. Journal of molecular biology 31, 507–518 (1968).

11. Boesen, T. O. et al. Dispensability of extrinsic DnaA regulators in Escherichia coli cell-cycle control. Proceedings of the National Academy of Sciences 121, (2024).

12. Zheng, H. et al. General quantitative relations linking cell growth and the cell cycle in Escherichia coli. Nature Microbiology 2020 5:8 5, 995–1001 (2020).

13. Katayama, T., Kasho, K. & Kawakami, H. The DnaA Cycle in Escherichia coli: Activation, Function and Inactivation of the Initiator Protein. Frontiers in Microbiology 8, (2017).

14. Fujimitsu, K., Senriuchi, T. & Katayama, T. Specific genomic sequences of E. coli promote replicational initiation by directly reactivating ADP-DnaA. Genes & Development 23, 1221–1233 (2009).

15. Kasho, K. & Katayama, T. DnaA binding locus datA promotes DnaA-ATP hydrolysis to enable cell cycle-coordinated replication initiation. Proceedings of the National Academy of Sciences 110, 936–941 (2013).

16. Kato, J. I. & Katayama, T. Hda, a novel DnaA-related protein, regulates the replication cycle in Escherichia coli. The EMBO Journal 20, 4253–4262 (2001).

17. Sekimizu, K. & Kornberg, A. Cardiolipin activation of dnaA protein, the initiation protein of replication in Escherichia coli. Journal of Biological Chemistry 263, 7131–7135 (1988).

18. Rozgaja, T. A. et al. Two oppositely oriented arrays of low-affinity recognition sites in oriC guide progressive binding of DnaA during Escherichia coli pre-RC assembly. Molecular Microbiology 82, 475–488 (2011).

19. Grimwade, J. E. et al. Origin recognition is the predominant role for DnaA-ATP in initiation of chromosome replication. Nucleic Acids Research 46, 6140–6151 (2018).

20. Ozaki, S. et al. A common mechanism for the ATP-DnaA-dependent formation of open complexes at the replication origin. Journal of Biological Chemistry 283, 8351–8362 (2008).

21. Hill, N. S., Kadoya, R., Chattoraj, D. K. & Levin, P. A. Cell size and the initiation of DNA replication in bacteria. PLoS Genet 8, e1002549 (2012).

22. Sompayrac, L. et al. Autorepressor Model for Control of DNA Replication. Nature New Biology 241, 133– 135 (1973).

23. Hansen, F. G., Christensen, B. B. & Atlung, T. The initiator titration model: computer simulation of chromosome and minichromosome control. Research in Microbiology 142, 161–167 (1991).

24. Olivi, L. et al. The Escherichia coli replication initiator DnaA is titrated on the chromosome. Nat Commun 16, 7813 (2025).

25. Berger, M. & ten Wolde, P. R. Robust replication initiation from coupled homeostatic mechanisms. Nature Communications 2022 13:1 13, 1–13 (2022).

26. Fu, H., Xiao, F. & Jun, S. Bacterial Replication Initiation as Precision Control by Protein Counting. PRX Life 1, 013011 (2023).

27. Iuliani, I., Mbemba, G., Lagomarsino, M. C. & Sclavi, B. Direct single-cell observation of a key Escherichia coli cell-cycle oscillator. Science Advances 10, (2024).

28. Katayama, T. The mutant DnaAcos protein which overinitiates replication of the Escherichia coli chromosome is inert to negative regulation for initiation. Journal of Biological Chemistry 269, 22075– 22079 (1994).

29. Carr, K. M. & Kaguni, J. M. The A184V missense mutation of the dnaA5 and dnaA46 alleles confers a defect in ATP binding and thermolability in initiation of Escherichia coli DNA replication. Molecular microbiology 20, 1307–1318 (1996).

30. Katayama, T., Crooke, E. & Sekimizu, K. Characterization of Escherichia coli DnaAcos protein in replication systems reconstituted with highly purified proteins. Molecular Microbiology 18, 813–820 (1995).

31. Klumpp, S., Zhang, Z. & Hwa, T. Growth Rate-Dependent Global Effects on Gene Expression in Bacteria. Cell 139, 1366–1375 (2009).

32. Campbell, J. L. & Kleckner, N. E. coli oriC and the dnaA gene promoter are sequestered from dam methyltransferase following the passage of the chromosomal replication fork. Cell 62, 967–979 (1990).

33. Waldminghaus, T. & Skarstad, K. The Escherichia coli SeqA protein. Plasmid 61, 141–150 (2009).

34. Fujimitsu, K. et al. Modes of Overinitiation, dnaA Gene Expression, and Inhibition of Cell Division in a Novel Cold-Sensitive hda Mutant of Escherichia coli. J Bacteriol 190, 5368–5381 (2008).

35. Charbon, G. et al. Suppressors of DnaA-ATP imposed overinitiation in Escherichia coli. Molecular Microbiology 79, 914–928 (2011).

36. Charbon, G. et al. Re-wiring of energy metabolism promotes viability during hyperreplication stress in E. coli. 13, e1006590 (2017).

37. Saluja, D. & Godson, G. N. Biochemical characterization of Escherichia coli temperature-sensitive dnaB mutants dnaB8, dnaB252, dnaB70, dnaB43, and dnaB454. J Bacteriol 177, 1104–1111 (1995).

38. Flåtten, I., Fossum-Raunehaug, S., Taipale, R., Martinsen, S. & Skarstad, K. The DnaA Protein Is Not the Limiting Factor for Initiation of Replication in Escherichia coli. PLOS Genetics 11, e1005276 (2015).

39. Von Freiesleben, U., Rasmussen, K. V., Atlung, T. & Hansen, F. G. Rifampicin-resistant initiation of chromosome replication from oriC in ihf mutants. Molecular Microbiology 37, 1087–1093 (2000).

40. Morigen Molina, F. & Skarstad, K. Deletion of the datA Site Does Not Affect Once-per-Cell-Cycle Timing but Induces Rifampin-Resistant Replication. Journal of Bacteriology 187, 3913–3920 (2005).

41. Olivi, L. et al. Live-cell imaging reveals the trade-off between target search flexibility and efficiency for Cas9 and Cas12a. Nucleic Acids Research 1, 13–14 (2024).

42. Zheng, H. et al. Interrogating the Escherichia coli cell cycle by cell dimension perturbations. Proceedings of the National Academy of Sciences 113, 15000–15005 (2016).

43. Berger, M. & ten Wolde, P. R. Synchronous Replication Initiation of Multiple Origins. PRX Life 1, 013007 (2023).

44. Li, D. et al. Extrusion-modulated DnaA activity oscillations coordinate DNA replication with biomass growth. eLife 14, RP107214 (2025).

45. Guimaraes, J. C., Rocha, M. & Arkin, A. P. Transcript level and sequence determinants of protein abundance and noise in Escherichia coli. Nucleic Acids Res 42, 4791–4799 (2014).

46. Lu, P., Vogel, C., Wang, R., Yao, X. & Marcotte, E. M. Absolute protein expression profiling estimates the relative contributions of transcriptional and translational regulation. Nat Biotechnol 25, 117–124 (2007).

47. LeBowitz, J. H. & McMacken, R. The Escherichia coli DnaB replication protein is a DNA helicase. Journal of Biological Chemistry 261, 4738–4748 (1986).

48. Gon, S. et al. A novel regulatory mechanism couples deoxyribonucleotide synthesis and DNA replication in Escherichia coli. EMBO J 25, 1137–1147 (2006).

49. Babu, V. M. P., Itsko, M., Baxter, J. C., Schaaper, R. M. & Sutton, M. D. Insufficient levels of the nrdAB-encoded ribonucleotide reductase underlie the severe growth defect of the Δhda E. coli strain. Mol Microbiol 104, 377–399 (2017).

50. Charbon, G., Bjørn, L., Mendoza-Chamizo, B., Frimodt-Møller, J. & Løbner-Olesen, A. Oxidative DNA damage is instrumental in hyperreplication stress-induced inviability of Escherichia coli. Nucleic Acids Res 42, 13228–13241 (2014).

51. Noguchi, Y. & Katayama, T. The Escherichia coli Cryptic Prophage Protein YfdR Binds to DnaA and Initiation of Chromosomal Replication Is Inhibited by Overexpression of the Gene Cluster yfdQ-yfdR-yfdS-yfdT. Front. Microbiol. 7, (2016).

52. Shinagawa, H., Iwasaki, H., Ishino, Y. & Nakata, A. SOS-inducible DNA polymerase II of E. coli is homologous to replicative DNA polymerase of eukaryotes. Biochimie 73, 433–435 (1991).

53. Napolitano, R., Janel-Bintz, R., Wagner, J. & Fuchs, R. P. P. All three SOS-inducible DNA polymerases (Pol II, Pol IV and Pol V) are involved in induced mutagenesis. The EMBO Journal 19, 6259–6265 (2000).

54. Truglio, J. J., Croteau, D. L., Van Houten, B. & Kisker, C. Prokaryotic Nucleotide Excision Repair: The UvrABC System. Chem. Rev. 106, 233–252 (2006).

55. Tang, M. et al. UmuD′_2_ C is an error-prone DNA polymerase, Escherichia coli pol V. Proc. Natl. Acad. Sci. U.S.A. 96, 8919–8924 (1999).

56. Reuven, N. B., Arad, G., Maor-Shoshani, A. & Livneh, Z. The Mutagenesis Protein UmuC Is a DNA Polymerase Activated by UmuD′, RecA, and SSB and Is Specialized for Translesion Replication. Journal of Biological Chemistry 274, 31763–31766 (1999).

57. Bell, J. C. & Kowalczykowski, S. C. RecA: Regulation and Mechanism of a Molecular Search Engine. Trends in Biochemical Sciences 41, 491–507 (2016).

58. Augustin, L. B., Jacobson, B. A. & Fuchs, J. A. Escherichia coli Fis and DnaA proteins bind specifically to the nrd promoter region and affect expression of an nrd-lac fusion. J Bacteriol 176, 378–387 (1994).

59. Stringer, A. M., Fitzgerald, D. M. & Wade, J. T. Mapping the Escherichia coli DnaA-binding landscape reveals a preference for binding pairs of closely spaced DNA sites. Microbiology (Reading) 170, 001474 (2024).

60. Torrents, E. et al. NrdR Controls Differential Expression of the Escherichia coli Ribonucleotide Reductase Genes. J Bacteriol 189, 5012–5021 (2007).

61. Knöppel, A., Broström, O., Gras, K., Elf, J. & Fange, D. Regulatory elements coordinating initiation of chromosome replication to the Escherichia coli cell cycle. Proceedings of the National Academy of Sciences 120, e2213795120 (2023).

62. Wang, P. et al. Robust Growth of Escherichia coli. Current Biology 20, 1099–1103 (2010).

63. Olivi, L. et al. Towards a synthetic cell cycle. Nature Communications 2021 12:1 12, 1–11 (2021).

64. Willis, L. & Huang, K. C. Sizing up the bacterial cell cycle. Nature Reviews Microbiology 2017 15:10 15, 606– 620 (2017).

65. Gaut, N. J. & Adamala, K. P. Reconstituting Natural Cell Elements in Synthetic Cells. Advanced Biology 5, 2000188 (2021).

66. Salas, M., Holguera, I., Redrejo-Rodríguez, M. & Vega, M. de. DNA-binding proteins essential for protein-primed bacteriophage ϕ29 DNA replication. Frontiers in Molecular Biosciences 3, 37 (2016).

67. Sokolik, C. G., Bar-Dolev, M., Milo, R., Adamala, K. P. & Levy, M. Synthetic cells by the numbers. iScience 28, (2025).

68. Koster, C. C. et al. De novo design of synthetic microbial genomes. Nat Rev Bioeng 1–18 (2026) doi:10.1038/s44222-026-00410-0.

69. Cooper, G. M. The Eukaryotic Cell Cycle. in The Cell: A Molecular Approach. 2nd edition (Sinauer Associates, Inc., 2000).

70. Datsenko, K. A. & Wanner, B. L. One-step inactivation of chromosomal genes in Escherichia coli K-12 using PCR products. Proceedings of the National Academy of Sciences of the United States of America 97, 6640– 6645 (2000).

71. Albert, H., Dale, E. C., Lee, E. & Ow, D. W. Site-specific integration of DNA into wild-type and mutant lox sites placed in the plant genome. The Plant journal: for cell and molecular biology 7, 649–659 (1995).

72. Creutzburg, S. C. A. Structure-function relations of RNA molecules involved in gene expression and host defence. https://doi.org/10.18174/532325 (2020) doi:10.18174/532325.

73. An Engineering Approach for Rewiring Microbial Metabolism. in Methods in Enzymology vol. 608 329–367 (Academic Press, 2018).

74. Doellinger, J., Schneider, A., Hoeller, M. & Lasch, P. Sample Preparation by Easy Extraction and Digestion (SPEED) - A Universal, Rapid, and Detergent-free Protocol for Proteomics Based on Acid Extraction. Mol Cell Proteomics 19, 209–222 (2020).

75. Abele, M. et al. Unified Workflow for the Rapid and In-Depth Characterization of Bacterial Proteomes. Mol Cell Proteomics 22, 100612 (2023).

76. Wu, C. et al. Enzyme expression kinetics by Escherichia coli during transition from rich to minimal media depends on proteome reserves. Nat Microbiol 8, 347–359 (2023).

77. Demichev, V., Messner, C. B., Vernardis, S. I., Lilley, K. S. & Ralser, M. DIA-NN: neural networks and interference correction enable deep proteome coverage in high throughput. Nat Methods 17, 41–44 (2020).

78. Gerault, M.-A., Camoin, L. & Granjeaud, S. DIAgui: a Shiny application to process the output from DIA-NN. Bioinformatics Advances 4, vbae001 (2024).

79. Schwanhäusser, B. et al. Global quantification of mammalian gene expression control. Nature 473, 337– 342 (2011).

80. Cox, J. et al. Accurate proteome-wide label-free quantification by delayed normalization and maximal peptide ratio extraction, termed MaxLFQ. Mol Cell Proteomics 13, 2513–2526 (2014).

81. Meng, C. Fast analyzing, exploring and sharing quantitative omics data using omicsViewer. 2022.03.10.483845 Preprint at 10.1101/2022.03.10.483845 (2022).

82. Cutler, K. J. et al. Omnipose: a high-precision morphology-independent solution for bacterial cell segmentation. Nat Methods 19, 1438–1448 (2022).

83. Bremer, H. & Churchward, G. An examination of the Cooper-Helmstetter theory of DNA replication in bacteria and its underlying assumptions. J Theor Biol 69, 645–654 (1977).

84. Ferullo, D. J., Cooper, D. L., Moore, H. R. & Lovett, S. T. Cell cycle synchronization of Escherichia coli using the stringent response, with fluorescence labeling assays for DNA content and replication. Methods 48, 8– 13 (2009).

