## Supplementary Information for "Establishment of titration-based control of DNA replication in *Escherichia coli*"

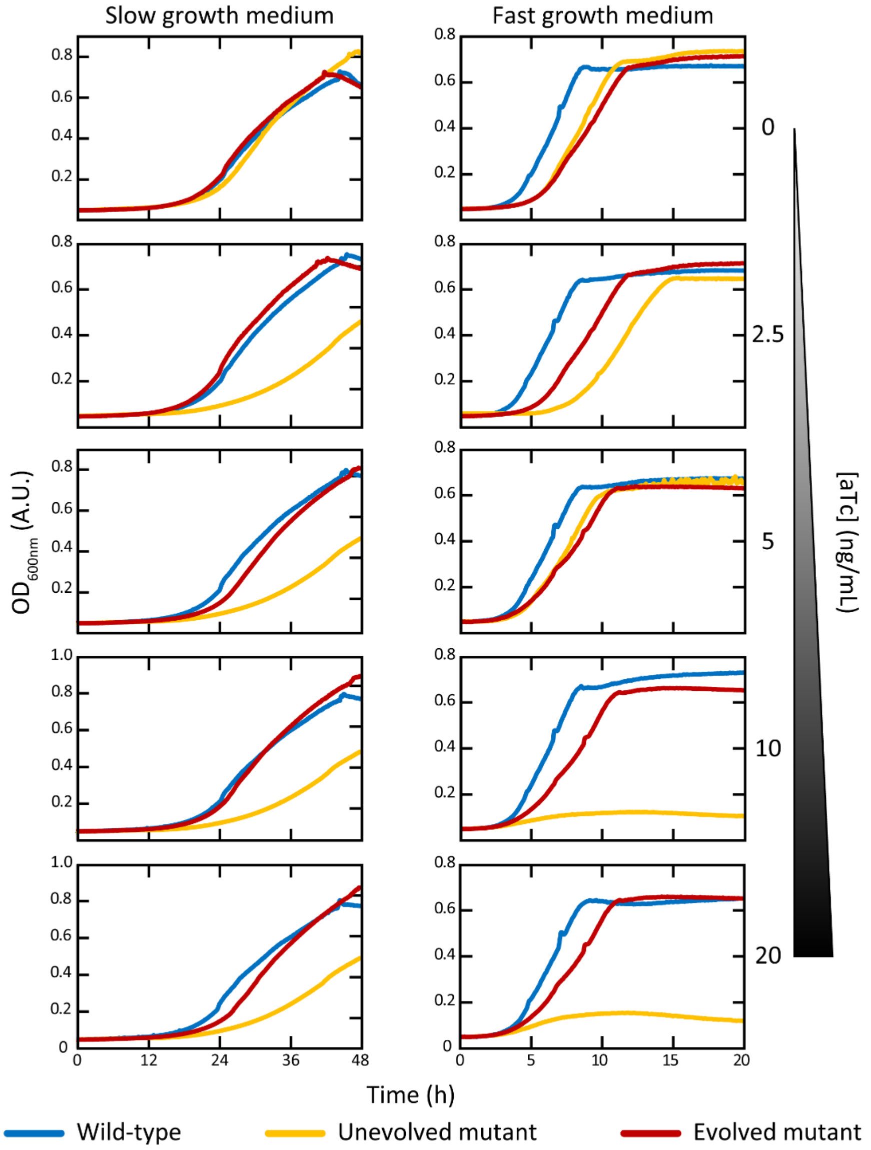


**Figure S1. Growth phenotype of *E. coli* mutant strains.** Growth characterisation of the wild-type *E. coli* MG1655 (blue line), the original mutant strain carrying the $p_{LtetO\text{-}1}$-*tetR-dnaAcos* locus (yellow line) and the mutant strain after evolution in fast growth and 10 ng/mL of aTc (red line). Cells were grown in either slow growth (left) or fast growth medium (right), each supplemented with five concentrations of aTc (0, 2.5, 5, 10 and 20 ng/mL).

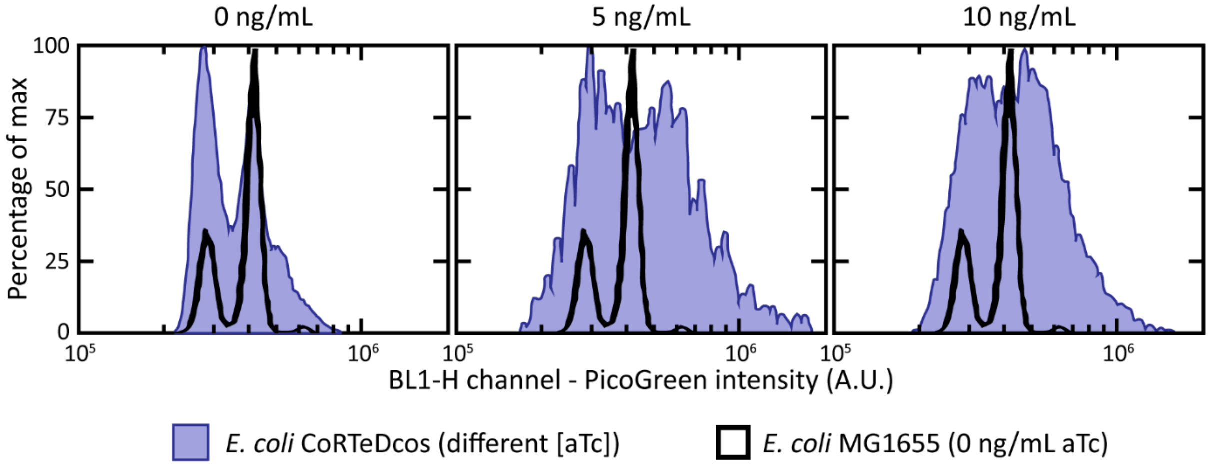


**Figure S2. *E. coli* CoRTeDcos is resistant to rifampicin treatment.** To obtain a measure of the average number of origins of replication in a *E. coli* population, cells are treated with rifampicin and cephalexin for at least 6 hours. This treatment stops new initiation events, allows any ongoing replication round to finish and prevents cell division. As a result, population of wild-type *E. coli* cells normally arrange in discrete peaks, with each peak indicating a integer increment in number of origins (black lines). On the contrary, *E. coli* CoRTeDcos shows a less defined division in populations even at the lowest level of induction (no aTc present, left). Whereas there was a increase in DNA content between conditions, the histograms did not represent synchronised populations were cells stopped replicating in the time frame of antibiotic treatment.

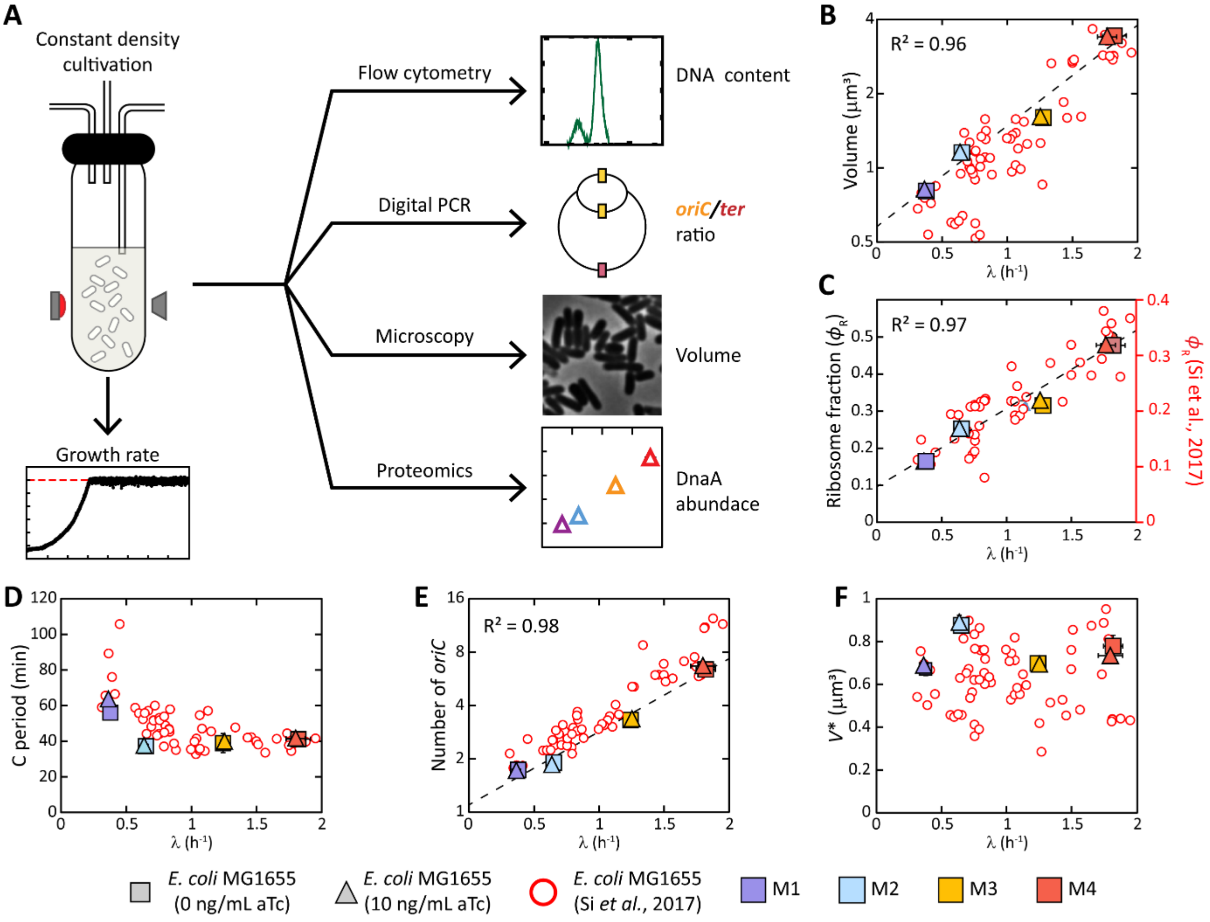


**Figure S3. Validation of our cultivation approach. A)** In our set-up, cell growth is balanced in exponential phase inside turbidostat reactors, maintaining the turbidity of the cell culture at a constant $OD_{600nm}$ = 0.3. After 10 generations, cells are harvested in different aliquots to be subjected to either flow cytometry for DNA content measurements, digital PCR for determination of *oriC/ter* ratio, microscopy for volume measurements or quantitative proteomics. We validated our cultivation strategy by recreating quantitative behaviours of *E. coli,* such as **B)** increase of volume with growth rate, **C)** increase of proteome ribosome fraction over growth rate, **D)** stable C period times, **E)** increase of the number of origins of replication over growth rate and **F)** stable volumes of initiation over growth rate. Data points are average of replicates and standard deviations. Data from *(1)* are plotted for comparison.

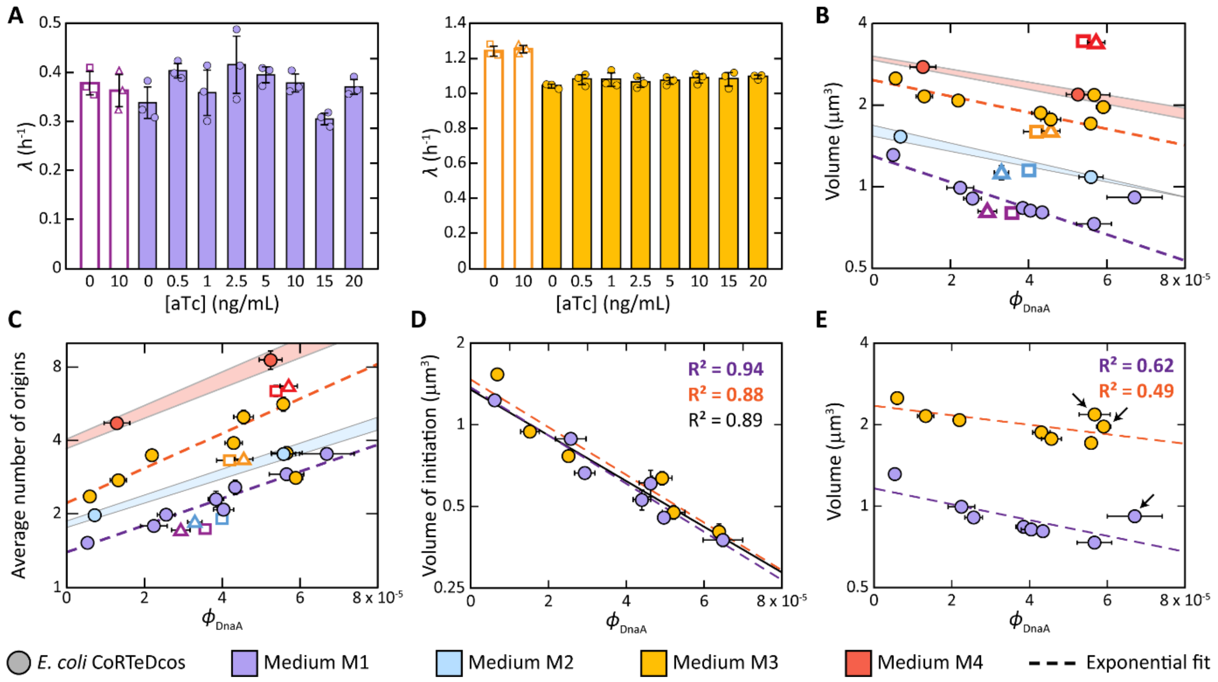


**Figure S4. Inducer-dependent behaviour of *E. coli* CoRTeDcos. A)** Different levels of inducer do not impact growth rate in *E. coli* CoRTeDcos in medium M1 and M3. See Figure 2 in the main text for the effect of aTc on the growth of *E. coli* CoRTeDcos in medium M2 and M4. **B)** Volume and **C)** number of origins of replication of *E. coli* CoRTeDcos at different DnaA mass fraction ($\phi_{DnaA}$) in all media. For media M1 and M3, the exponential relationships are the same as in the main text (Figure 3B). For media M2 and M4, the shaded regions represent hypothetical trends, based on the available measurements and their standard deviations. **D)** Individual exponential relationships between $\phi_{DnaA}$ and volume of initiation in media M1 and M3. The exponential relationship derived from *E. coli* CoRTeDcos in all media and inducer concentrations (black continuous line) is the same as in the main text (Figure 3D). **D)** Trendlines of the relationship between $\phi_{DnaA}$ and volume for *E. coli* CoRTeDcos in media M1 and M3 including measurements coming from all inducer concentrations. Measurements from cell grown in medium M1 with 20 ng/mL aTc and in medium M3 with either 15 or 20 ng/mL aTc (indicated by arrows) deviate from the general exponential relationship. They were considered as outliers and excluded from the exponential fits of the main text. Purple, blue, yellow and red symbols represent data acquired from cells grown in media M1, M2, M3 and M4, respectively. All data points are the average of three independent biological replicates and error bars represent the standard deviation. In some cases, error bars are smaller than the symbols. In **A)**, individual data points are plotted.

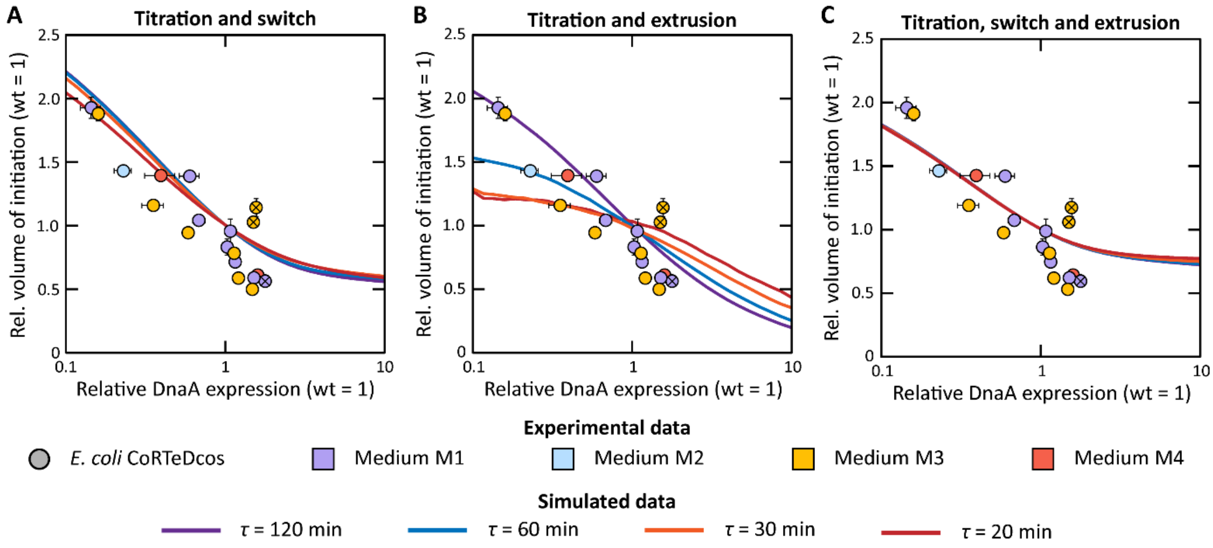


**Figure S5. Comparison of experimental *E. coli* CoRTeDcos data with different models of *E. coli* DNA replication initiation.** The experimental relationship between the relative expression of DnaA and the relative volume of initiation (normalised for the average of all values in the same medium) is compared with prediction coming from simulations of *E. coli* DNA replication using different models: **A)** titration of DnaA and interconversion between DnaA-ATP/DnaA-ADP ($\chi_\nu^2 \approx 45$); **B)** titration of DnaA and extrusion modulated by the nucleoid-associated protein H-NS ($\chi_\nu^2 \approx 71$); **C)** titration of DnaA, DnaA-ATP/DnaA-ADP interconversion and extrusion modulated by H-NS ($\chi_\nu^2 \approx 54$). Purple, blue, yellow and red symbols indicate experimental data collected in medium M1, M2, M3 and M4, respectively. Purple, blue, orange and red solid lines indicate simulated trends for cells with doubling times of 120 min, 60 min, 30 min and 20 min, respectively. All data points are the average of three independent biological replicates and error bars represent the standard deviation. In some cases, error bars are smaller than the symbols.

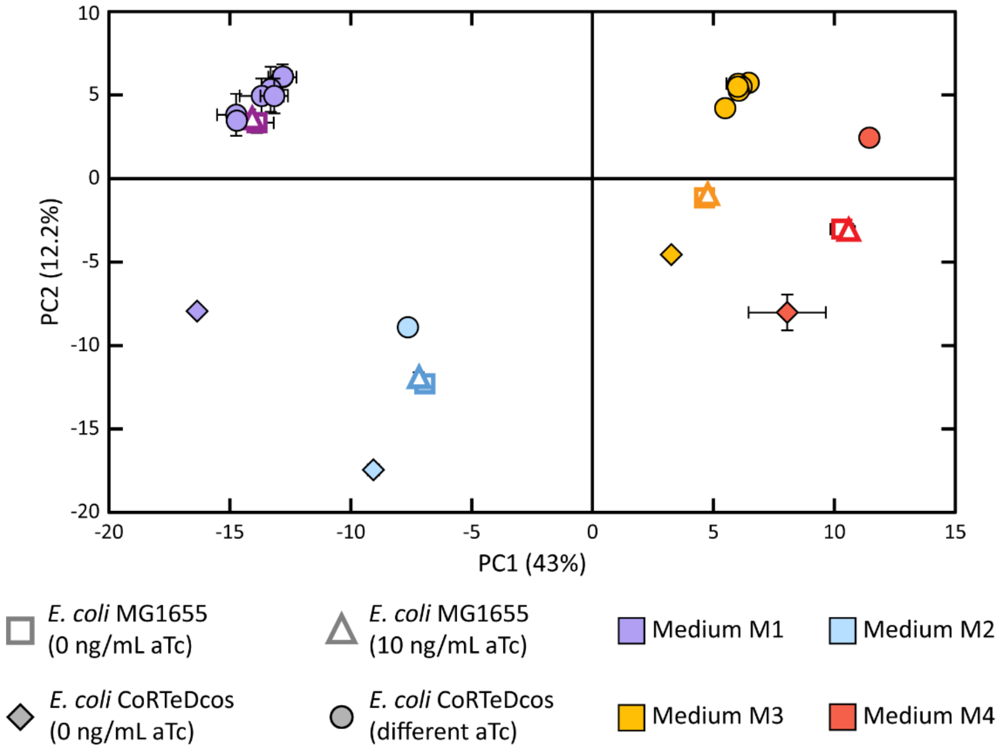


**Figure S6. Principal component analysis of the proteomes of *E. coli* MG1655 and *E. coli* CoRTeDcos in all the tested conditions.** In all growth condition, the proteome of *E. coli* CoRTeDcos only minorly changed in the presence of different concentrations of aTc (filled circles), but it was majorly altered in the absence of the inducer (filled diamonds). On the other hand, the proteome of *E. coli* MG1655 remained unchanged in either absence (empty squares) or presence of aTc (empty triangles), indicating that all of the changes observed in *E. coli* CoRTeDcos come from the different expression levels of DnaAcos.

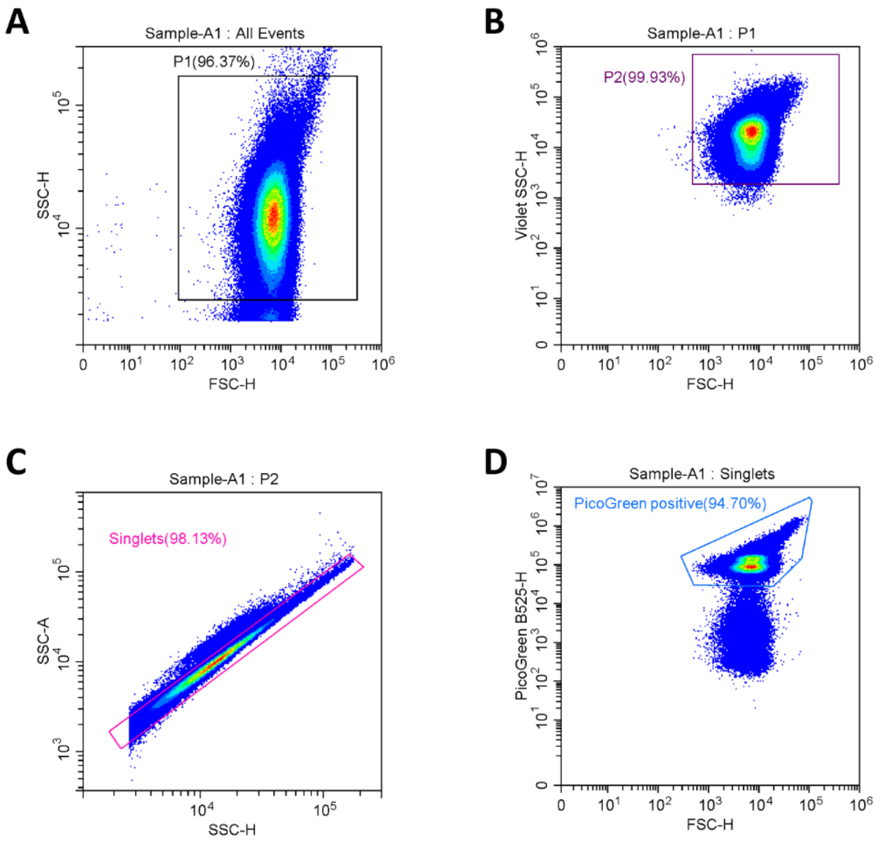


**Figure S7. Gating strategy employed in the measurement of DNA content. A)** The population of cells are gated from noise by plotting SSC-H against FSC-H using a blue laser (population P1). **B)** Cells are then more finely gated from noise by plotting the SSC-H obtained using a violet laser versus the same FSC-H (population P2). **C)** For population P2, SSC-H is then plotted against SSC-A and the Singlets population is gated from doublet events by gating on the diagonal. **D)** Finally, the intensity of the PicoGreen dye in the Singlets population is obtained and the positive cells for PicoGreen are gated from the rest. Histograms of DNA contents (see Figure S2 for an example) are then obtained from the PicoGreen positive population.

**Table S1. List of bacterial strains used in this study, with related genotypes and source.**

| Strain | Characteristics | Source |
|---|---|---|
| *E. coli* MG1655 | K-12; $F^-$ $\lambda^-$ *rph-1* | ATCC |
| *E. coli* DH5α | MG1655; *endA1 glnV44 thi-1 recA1 relA1 gyrA96 deoR nupG* ϕ80d*lacZ*ΔM15 Δ(*lacZYA-argF)U169 hsdR17* ($r_K^-$ $m_K^+$) | New England Biolabs |
| *E. coli* DH5α x pCoRe-T | DH5α harbouring the pCoRe-T plasmid | This study |
| *E. coli* DH5α x pCoRe-Dcos | DH5α harbouring the pCoRe-Dcos plasmid | This study |
| *E. coli* pre-CoRTeDcos | MG1655; Δ*dnaAp-dnaA*::lox66/71-*pLtetO-1-tetR-dnaAcos*-lox66/71 | This study |
| *E. coli* CoRTeDcos | Pre-CoRTeDcos; *dnaB*(G726>T) | This study |

**Table S2. Complete chemical composition of cultivation media used this study.**

| Identifier | Buffer/base composition | Carbon source | Supplement |
|---|---|---|---|
| Medium M1 | $K_2HPO_4$ 36.7 mM, NaCl 8.6 mM, $Na_2HPO_4$ 47.8 mM, $NH_4Cl$ 18.7 mM, $CaCl_2$ 0.5 mM, $MgCl_2$ 1 mM, $FeCl_3$ 13 µM, $ZnCl_2$ 6.2 µM, $CuCl_2$ 0.76 µM, $CoCl_2$ 0.42 µM, $H_3BO_3$ 1.62 µM, $MnCl_2$ 0.081 µM | Glycerol 22 mM | / |
| Medium M2 | | Glucose 22 mM | Leu 0.6 mM, Ile 0.6 mM, Val 0.6 mM, thiamine 0.03 mM |
| Medium M3 | | Glucose 22 mM | Arg 1.15 mM, Asn 0.38 mM, Asp 0.15 mM, Cys 0.21 mM, Glu 0.14 mM, Gly 0.13 mM, His 0.1 mM, Ile 0.38 mM, Leu 0.38 mM, Lys 0.22 mM, Met 0.1 mM, Phe 0.09 mM, Pro 0.17 mM, Ser 0.29 mM, Thr 0.17 mM, Trp 0.025 mM, Tyr 0.11 mM, Val 0.17 mM |
| Medium M4 | Pancreatic digest of casein 17 g/L, papaic digest of soybean 3 g/L, NaCl 85 mM, $K_2HPO_4$ 14 mM | Glucose 22 mM | / |

**Table S3. List of plasmids used in this study.**

| Plasmid | Characteristics | Source |
|---|---|---|
| pSC020 | *repA101*(Ts), *oriV, ampR, araC,* $P_{BAD}$*-lRed, lacIq,* $P_{Lac}$*-Cre* | 58 |
| pCoRe-T | pUC19 backbone; *ampR, lox66-cmR-lox71, pLtetO-1-tetR, BsaI* | This study |
| pCoRe-Dcos | pUC19 backbone; *ampR, BsaI, dnaAcos, lox66-kanR-lox71* | This study |

**Table S4. List of primers and chemically synthesised DNA used in this study.** Sequences introducing homology arms for recombination into the genome of *E. coli* MG1655 are highlighted in bold.

| Identifier | Sequence (5′-3′) | Used for |
|---|---|---|
| BG22166 | cgagtggagtccgccatgtcactttcgctttggcagcagtgtcttgcccgattgcagg<br>atgagttaccagccacagaattcagtatgtggatacgcccattgcaggcggaactg<br>agcgataacacgctggccctgtacgcgccaaaccgttttgtcctcgattgggtacgg<br>gacaagtaccttaataatatcaatggactgctaaccagtttctgcggagcggatgcc<br>ccacagctgcgttttgaagtcggcaccaaaccggtgacgcaaacgccacaagcgg<br>cagtgacgagcaacgtcgcggcccctgcacaggtggcgcaaacgcagccgcaac<br>gtgctgcgccttctacgcgctcaggttgggataacgtcccggccccggcagaaccg<br>acctatcgttctaacgtaaacgtcaaacacacgtttgataacttcgttgaaggtaaat<br>ctaaccaactggcgcgcgcggcggctcgccTTgtggcggataaccctggcggtgc<br>ctataacccgttgttcctttatggcggcacgggtctgggtaaaactcacctgctgcat<br>gTTgtgggtaacggcattatggcgcgcaagccgaatgccaaagtggtttatatgc<br>actccgagcgctttgttcaggacatggttaaagccctgcaaaacaacgcgatcgaa<br>gagtttaaacgctactaccgttccgtagatgcactgctgatcgacgatattcagttttt<br>tgctaataaagaacgatctcaggaagagtttttcTATaccttcaacgccctgctgga<br>aggtaatcaacagatcattctcacctcggatcgcCACccgaaagagatcaacggc<br>gttgaggatcgtttgaaatcccgcttcggttggggactgactgtggcgatcgaaccg<br>ccagagctggaaacccgtgtggcgatcctgatgaaaaaggccgacgaaaacga<br>cattcgtttgccgggcgaagtggcgttctttatcgccaagcgtctacgatctaacgta<br>cgtgagctggaaggggcgctgaaccgcgtcattgccaatgccaactttaccggac<br>gggcgatcaccatcgacttcgtgcgtgaggcgctgcgcgacttgctggcattgcag<br>gaaaaactggtcaccatcgacaatattcagaagacggtggcggagtactacaag<br>atcaaagtcgcggatctcctttccaagcgtcgatcccgctcggtggcgcgtccgcgc<br>cagatggcgatggcgctggcgaaagagctgactaaccacagtctgccggagatt<br>ggcgatgcgtttggtggccgtgaccacacgacggtgcttcatgcctgccgtaagatc<br>gagcagttgcgtgaagagagccacgatatcaaagaagatttttcaaatttaatcag<br>aacattgtcatcgtaaacctat | Synthetic DNA of *dnaAcos* |
| BG22660 | **gcctcgcggcaggatcgtttacacttagcgagttctggaaagtcctgtgg**gctagtt<br>attgctcagcggtgg | Amplified pCoRe-T fragment |
| BG20764 | gcgtttcggtgatgacggtgaaaac | Amplified pCoRe-T fragment |
| BG21406 | cgattcattaatgcagctggcacg | Amplified pCoRe-Dcos fragment |
| BG22661 | **taaatgctcacgttctacggtaaatttcataggtttacgatgacaatgtt**ccgactgg<br>aaagctaccg | Amplified pCoRe-Dcos fragment |
| BG22467 | cggcaatttccgcgccttc | Amplified *dnaA* locus in *E. coli* genome |
| BG22496 | gacctcacaccagtggaaaccag | Amplified *dnaA* locus in *E. coli* genome |
| SJO416 | ttcaccagatgggcaatcacttcgg | Amplified *oriC* in digital PCR |
| SJO417 | acattgtggtggtgcataacggcatc | Amplified *oriC* in digital PCR |
| SJO432 | atggatcttcctgctataccgccac | Amplified *ter* in digital PCR |
| SJO433 | ggccaacatttgcgtccacagataag | Amplified *ter* in digital PCR |

**Table S5. Whole genome sequencing results of the strain used in this study.** NC_000913.3 is the accession number of *E. coli* K-12 MG1655 reference genome.

| Strain | Reference genome | Mutations compared to reference |
|---|---|---|
| *E. coli* MG1655 | NC_000913.3 | 276’697 C→A (*mmuP* A458E); 1’873’031 IS1(–) + 9 bp (*dgcJ*); 4’296’381 IS +GC (intergenic) |
| *E. coli* pre-CoRTeDcos | MG1655 | Δ*dnaAp-dnaA*::lox66/71-*pLtetO-1-tetR-dnaAcos*-lox66/71 |
| *E. coli* CoRTeDcos | Pre-CoRTeDcos | 4’265’708 G→T (*dnaB* M424I) |

**Table S6. Number of cells imaged per strain, condition and replicate to obtain cell size measurements.** In the "Total number of cells" column, the three numbers between brackets represent the number of cells imaged in replicate 1, replicate 2 and replicate 3, respectively.

| Strain | Medium | aTc (ng/mL) | Number of cells (replicates) |
|---|---|---|---|
| *E. coli* MG1655 | M1 | 0 | 6227 (2213, 2387, 1627) |
| | | 10 | 6581 (2381, 2031, 2169) |
| | M2 | 0 | 7408 (2498, 2370, 2540) |
| | | 10 | 7448 (2947, 2245, 2256) |
| | M3 | 0 | 6674 (2592, 2067, 2015) |
| | | 10 | 6779 (2286, 2399, 2094) |
| | M4 | 0 | 6965 (2739, 2284, 1942) |
| | | 10 | 6991 (2609, 2406, 1976) |
| *E. coli* CoRTeDcos | M1 | 0 | 6313 (2097, 2121, 2095) |
| | | 0.5 | 7017 (2621, 2398, 1998) |
| | | 1 | 7108 (2588, 2453, 2067) |
| | | 2.5 | 6403 (2120, 2250, 2033) |
| | | 5 | 6582 (2150, 2036, 2396) |
| | | 10 | 6990 (2767, 1997, 2226) |
| | | 15 | 7057 (2118, 2094, 2845) |
| | | 20 | 6888 (2353, 2144, 2391) |
| | M2 | 0 | 4550 (1390, 1857, 1303) |
| | | 10 | 6740 (2063, 2325, 2352) |
| | M3 | 0 | 6611 (2852, 2205, 1554) |
| | | 0.5 | 6053 (2022, 2135, 1896) |
| | | 1 | 6385 (2316, 2096, 1973) |
| | | 2.5 | 7983 (2714, 2706, 2563) |
| | | 5 | 6496 (2207, 2427, 1862) |
| | | 10 | 6007 (1794, 2210, 2003) |
| | | 15 | 6530 (2659, 2305, 1566) |
| | | 20 | 6328 (2455, 1742, 2131) |
| | M4 | 0 | 7035 (2389, 2575, 2071) |
| | | 10 | 6906 (2493, 2173, 2240) |

**Table S7. Parameters used in the cell cycle simulations.** With the exception of doubling time, all other parameters were left as in the original work[44] to recreate the established relationships.

| Parameter | Values |
|---|---|
| Doubling time ($\tau$) | 20 min, 30 min, 60 min, 120 min |
| DnaA synthesis rate ($\alpha_A$) | From 60 $h^{-1}$ $\mu m^{-3}$ to 3840 $h^{-1}$ $\mu m^{-3}$ |
| *Used in titration and titration-extrusion models* | |
| Critical DnaA free concentration ($[A_f^c]$) | 10 $\mu m^{-3}$ |
| Extruder synthesis rate ($\alpha_H$) | 180 $h^{-1}$ $\mu m^{-3}$ |
| *Used in titration-switch and titration-switch-extrusion models* | |
| Total DnaA concentration ($[D]_T$) | 400 $\mu m^{-3}$ |
| Lipids activation rate ($\alpha_l$) | 750 $h^{-1}$ |
| *DARS1* activation rate ($\alpha_{d1}$) | 100 $h^{-1}$ |
| *DARS2* activation rate ($\alpha_{d2}$) | 643 $h^{-1}$ |
| *datA* deactivation rate ($\beta_{datA}$) | 600 $h^{-1}$ |
| RIDA deactivation rate ($\beta_{RIDA}$) | 500 $h^{-1}$ |
| Dissociation constant of DnaA activation/deactivation ($K_D$) | 50 $\mu m^{-3}$ |
| Dissociation constant of DnaA promoter ($K_D^P$) | 400 $\mu m^{-3}$ |
| Cooperativity of *dnaA* expression ($n$) | 5 |
| Extruder synthesis rate ($\alpha_H$) | 550 h-1 $\mu m^{-3}$ |
| Critical DnaA-ATP concentration ($[A_{ATP}^{fc}]$) | 200 $\mu m^{-3}$ |
